# Why Many Molecular Simulation Research Findings Might Be False: An Analysis of Inter-Simulations Differences Based on Simulation Time and Number of Replicas

**DOI:** 10.1101/2022.08.23.504950

**Authors:** Bernhard Knapp, Charlotte M. Deane

## Abstract

Molecular simulations are a common technique to investigate the dynamics of proteins, DNA and RNA. A typical application is the simulation of a wild-type structure and a mutant structure where the mutant has a significantly higher (or lower) potency to trigger a signalling cascade. The study would then analyse the observed differences between the wild-type and mutant simulation and link these to their differences. However differences in the simulations cannot always be reproduced by other research groups even if the same parameters as in the original simulations are used. This is caused by the rugged energy landscape of many biological structures which means that minor differences in hardware or software can cause simulation to take different paths. This would not be a problem if the simulation time would be infinitely long but in real life the simulation time is always finite.

In this study we use large scale molecular simulations of four different systems (a 10-mer peptide wild-type and mutant as well as a T-cell receptor, peptide and MHC complex as wild-type and mutant) with 100 replicas each totalling 620 000 ns to quantify the magnitude of (non-) reproducibility when comparing inter-simulation differences (e.g. wild-type vs mutant).

Using a bootstrapping approach we found that simulation times of at least 2 to 3 times the experimental folding time using a minimum of 3 replicas are necessary for reproducible results. However, for most complexes of interest such long simulation times are far out of reach which means that it is only possible to sample the local phase space neighbourhood of the x-ray structure. To sample this neighbourhood reliably around 10 to 20 replicas are needed.

**Graphical abstract:** 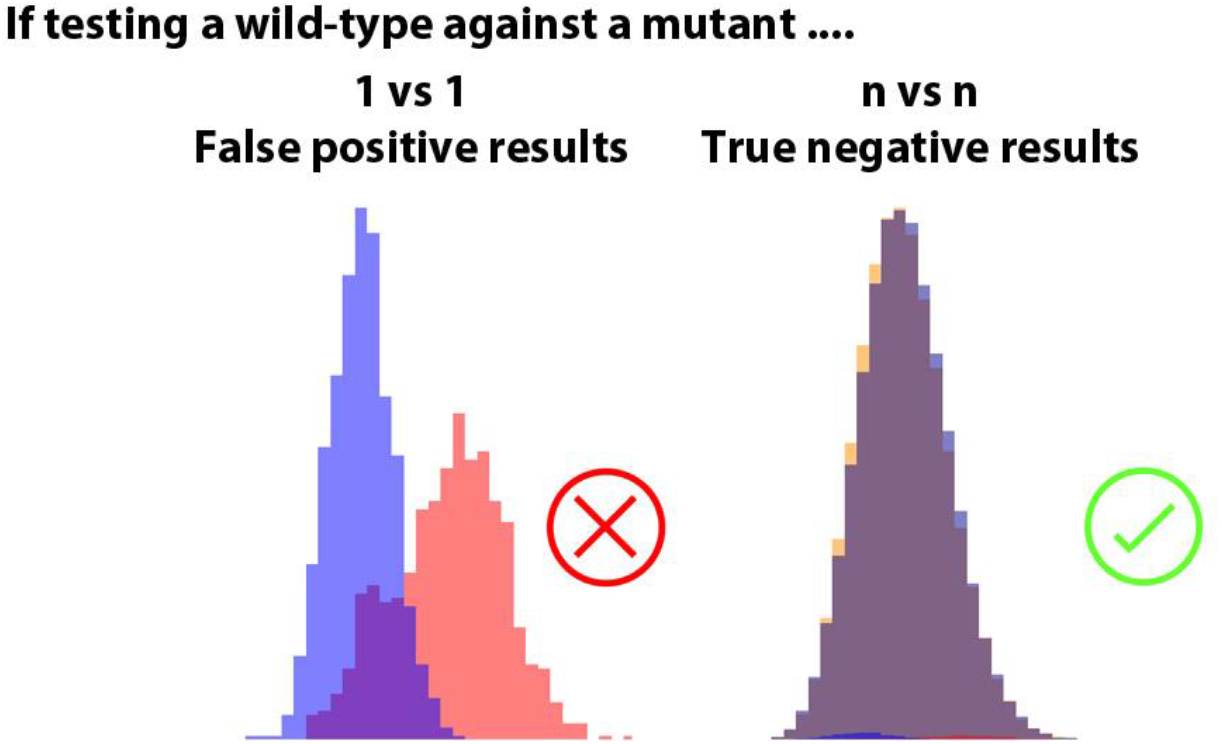

## Introduction

Molecular simulations are a widely used technique to investigate the dynamics and folding of proteins, DNA, and RNA ^1,2^. Newton’s equations of motions are solved for a given system of atoms over time. This allows insight into the dynamics of a system beyond the static image of an X-ray structure or computational model structure.

The energy landscape of molecular simulations is complex and often rugged containing a large number of local minima separated by high energy barriers ^3^. Running identical simulations multiple times should in theory give identical results but slight differences in underlying hardware, compiler optimisation levels, library versions, dynamic load balancing, parallelisation, system specific random number generators, or initial forces created randomly according to a Maxwell distribution can lead to differing trajectories ^4,5^. If a simulation runs close to an energy cliff one simulation might “fall” off the cliff while another might just about avoid it. From this moment on the two identical simulations will sample completely different areas of the energy landscape (solution/phase space). The interpretation of these results might lead two very different conclusions of the two identically parameterised simulations. This renders the question of how reproducible the results of molecular simulations are.

In the following we show how this issue can lead to misleading conclusions: An often seen study setup is to compare the simulation of a wild-type protein with the simulation of a mutant of the same protein. The wild-type and mutant are expected to develop two different trajectories on which basis the effect of the mutation can be described and understood. An example is shown in Figure 1. A T-cell receptor (TCR) in complex with a major histocompatibility complex (MHC) and the experimentally known immunogenic peptide FLRGRAYGL was simulated for 100 ns. A second simulation was performed using the identical parameters but before the single point mutation Y7A was introduced into the peptide. The Y7A mutation is known to dramatically impair cytotoxic T-cell recognition ^6^. From both simulations the number of hydrogen bonds (H-bonds) between the TCR α-chain and TCR β-chain were calculated (Figure 1A). It can be seen that the mutant simulation has on average at least 7 H-bonds less than the wild-type simulation which amounts to a loss of more than one third of the overall wild-type H-bonds. It could be concluded from these simulations that the mutation Y7A destabilises the TCR chain arrangement which is the underlying structural cause for the impaired T-cell recognition. This conclusion could be further strengthened by quoting the original X-ray structure paper ^6^ that states that Y7 of the peptide protrudes deeply between CDR1α and CDR3β into the TCR. However, there is one problem with this conclusion: It is wrong. If 100 replicas (identical simulations but different initial velocities) of both the wild-type and the Y7A mutant are performed and averaged then hardly any difference in the number of H-bonds is present (Figure 1B). The Y7A mutation does not disrupt the TCR chain arrangement and a 1vs1 molecular simulation study would have led to a false positive conclusion and publication.

**Figure 1:**
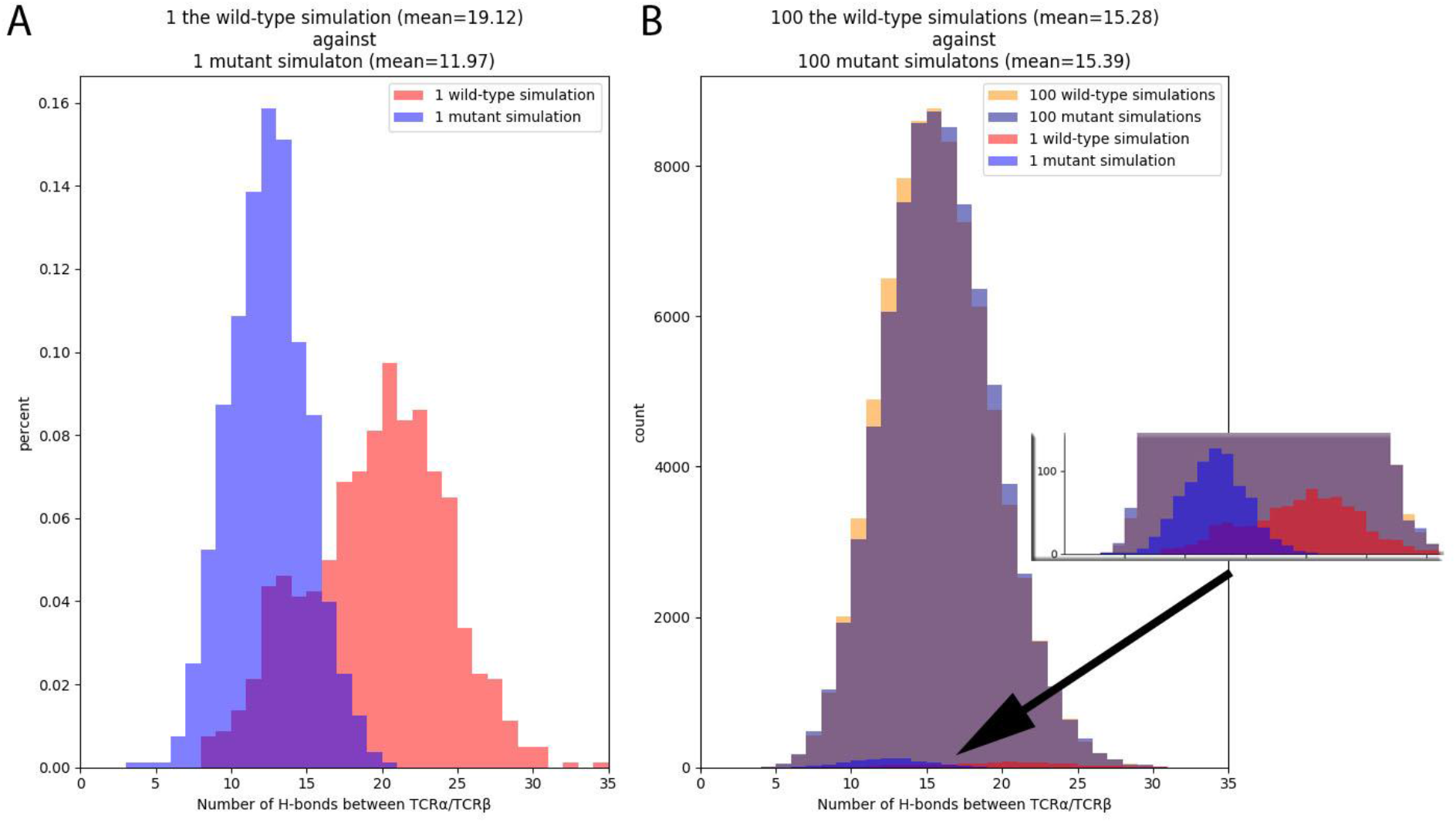
(A) Misleading 1vs1 comparison between a TCRpMHC simulation with an immunogenic wild-type peptide and a non-immunogenic Y7A mutant peptide that would lead to the conclusion that the mutant peptide destabilises the TCR chain arrangement. (B) Same as (A) but with 100 vs 100 replicas. The conclusion of (A) was wrong. The Y7A peptide mutation does not lower the number of H-bonds between the TCR chains. On average over 100 simulations of the wild-type and 100 simulations of the mutant the number of H-bonds between the TCR chains is almost identical. The inlay shows the two distributions of (A) relative to the distributions of all 200 replicas.

While it can relatively easily be shown that a 1vs1 comparison is prone to false positive conclusions by picking the most extreme examples as we have done in the example above the more interesting question is to quantify the average uncertainty and estimate the inaccuracy at a given number of replicas and simulation lengths.

In a recent study ^5^ we analysed this uncertainty for *intra*-simulation differences i.e. how different are replica simulations of the same structure to each other. In this study we go one step further and analyse *inter*-simulations differences. Firstly we compare simulations of two small peptides with each other as the phase space of such small systems can be explored in depth and complete sampling is likely to be achieved. Secondly we investigate the simulations of a wild-type TCRpMHC system against a mutant of the same system as described in the introduction.

On these grounds we numerically quantify the reproducibility and reliability of a wild-type vs mutant simulation setting. Based on our dataset it is possible to give a rough estimate on how likely it is that the results of a molecular simulation study can be reproduced given the simulations time and number of replicas.

## Methods

### Structures

#### Wild-type and mutant peptide systems

Firstl we used two peptide model systems as these are small enough to assume good coverage of their configuration spaces and thereby allow in-depth investigations of long term folding behaviour. We used the designed Chignolin 10-mer peptide GYDPETGTWG. This peptide was designed on the statistics of 10 000 protein segments and has despite its small size the elementary properties of a natural protein ^7^. The structure of this peptide is available from the Protein Data Bank (PDB) via accession code 1UAO ^7^ and is referred to as wild-type (WT) peptide.

For comparison we used a mutated version of Chignolin in which the C- and N-terminal Glycines were substituted with Tyrosines yielding the sequence YYDPETGTWY. This structure has even more protein-like characteristics, is therefore expected to be more stable and is available via PDB accession code 2RVD ^8^. It is referred to as mutant (MT) peptide. We investigated this MT in our previous intra-replica reproducibility study ^5^.

#### T-cell receptor and MHC systems

Secondly we used the Lc13 TCR binding the Epstein-Barr virus (EBV) 9-mer peptide FLRGRAYGL presented by the human MHC allele HLA-B*08:01 available via Protein Data Bank (PDB) accession code *1mi5* ^6^. This 826 amino acid structure is a good model system as it has been investigated intensively before ^6,9,10^ and our previous study ^5^ that investigated intra-replica variations used the same system. In all analyses the full TCR including the constant regions is used as we have recently shown that this is necessary for reliable results ^11^. For simulations with pMHC the whole pMHC including the α_3_-region and β_2_-microglobulin is used.

We created a total of three different starting structures: The wild-type TCRpMHC complex was directly taken from PDB accession code 1mi5. This system contains 826 amino acids and is from here on referred to as “TCRpMHC-WT”.

A mutant complex was created from PDB accession code 1mi5 by introducing the mutation Y7A into the 9-mer peptide. As described in the introduction this peptide position is known to be important for T-cell reactivity as it protrudes deeply into the TCR and fine specificity analysis of LC13 TCR cytotoxic T cell reactivity on peptide analogs has shown that most mutations of this position abrogate T-cell reactivity ^6^. To introduce the Y7A mutation we used SCWRL ^12^ via the peptX framework ^13^ as we have shown before that this tool yields the highest accuracy for peptide/MHC modelling ^14,15^. This structure contains also 826 amino acids and is referred to as “TCRpMHC-7A”.

#### Simulations

All simulations were run in GROMACS 4 ^16^ using the GROMOS 53a6 force field ^17^. The four structures were inserted into explicit dodecahedronic SPC3 ^18^ water baths that allowed for a minimum distance of 1.5 nm between protein/peptide and box boundary. Periodic boundary conditions were applied. Random water molecules were replaced with Na+ and Cl- ions to achieve a neutral charge as well as a salt concentration of 0.15 mol/liter. The systems were energetically minimised using the steepest descent algorithm and then warmed up to 310 K using position restraints. Each simulation of all four structures was carried out 100 times using different random velocities generated at 310 K according to a Maxwell distribution. This ensures that all 100 replicas have a slightly different start and it emulates the causes of differing trajectories discussed in the introduction right from the start of each simulation.

For the peptide WT and peptide MT the production run of each of the 100 replicas was 3000 ns long. This corresponds to about 500 the folding time of the WT and MT and it is likely that excellent coverage of the folding space is achieved. The MT simulations where taken from our previous study about intra-replica differences ^5^.

For the TCRpMHC-WT and TCRpMHC-7A systems production runs of 100 replicas of 100 ns each were carried out. This simulation length of 100 ns is a state-of-the-art for TCRpMHC simulations ^19^. The TCRpMHC-WT simulations were taken from previous studies of our group ^9,20–22^.

All simulations were carried out on the ARCUS-B cluster of the Oxford Advanced Research Computing (ARC) facility. An overview of the simulations is given in Table 1.

**Table 1:** Overview of simulations.

| Setup | TCR | MHC | Peptide | Number of amino acids | Length of one simulation | N replicas | Total simulation time |
| --- | --- | --- | --- | --- | --- | --- | --- |
| **TCRpMHC-WT** | LC13 | HLA-B*08 | FLRGRAYGL | 826 | 100 ns | 100 | 10 000 ns |
| **TCRpMHC-7A** | LC13 | HLA-B*08 | FLRGRAAGL | 826 | 100 ns | 100 | 10 000 ns |
|  | **Peptide** |  |  |  |  |  |  |
| **WT peptide** | GYDPETGTWG |  |  | 10 | 3 000 ns | 100 | 300 000 ns |
| **MT peptide** | YYDPETGTWY |  |  | 10 | 3 000 ns | 100 | 300 000 ns |
|  |  |  |  |  |  | Total: | 620 000 ns |

#### Trajectory analysis

Trajectories were evaluated using the Gromacs built-in functions gmx distance, hbond, gyration, rms and sas ^16^. Manual inspections of the simulations trajectories were done using VMD ^23^ and the vmdICE plugin ^24^ as well as the gro2mat package ^25^ and pyHVis3D ^26^.

#### Reproducibility analysis: bootstrapping

In order to determine the reliability and reproducibility of a specific difference between two sets of simulations (given a specific number of replicas and simulations length) we performed bootstrapping analyses. For example for the difference in the number of H-bonds between TCRpMHC-WT and TCRpMHC-7A we performed the following analysis: from the 100 TCRpMHC-WT replicas we sampled uniformly at random and with replacement n replicas. Then we did the same from the 100 TCRpMHC-7A replicas. For each of the two samples, we computed the average of the number of H-bonds over all frames and replicas. We repeated this sampling procedure 5000 times, and recorded the absolute value of the difference d_k_, k=1,2, …, 5000 between the TCRpMHC-WT sample and TCRpMHC-7A sample in each repetition. From this we computed the average over all 5000 d_k_ values. This average is shown as one point in the C subfigures of the following figures. In these figures each point represents a unique combination of number of replicas used (illustrated by colour coding) and length of the trajectory (which is artificially cut off at different time points (x-axis)). This procedure allows us to illustrate the influence of the number of replicas and the simulation time on the average difference between the two groups of simulations (e.g. TCRpMHC-WT vs TCRpMHC-7A).

#### Difference and Cohen’s d

In order to quantify the difference between two sets of simulations we use the simple difference between the mean values as given by:

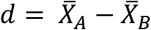

where 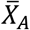 and 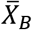 are the absolute mean values over all frames and all replicas of descriptor X (e.g. distance or radius or gyration of a specific area). A and B are the two sets of simulations to be compared (e.g. TCRpMHC-WT and TCRpMHC-7A).

As a second measure of difference we use the Cohen’s d effect size:

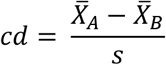

Where s is the pooled variance of 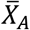 and 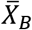 An effect size of ≥0.2 is considered a small effect, ≥0.5 a medium effect and ≥0.8 a large effect ^27^.

## Results

### Results of the peptide systems

First we describe the differences between the small peptide wild-type and mutant systems based on 200 peptide simulations (100 GYDPETGTWG and 100 YYDPETGTWY) of 3000 ns each.

### Distance between residue 1 and 10 of WT and MT

The distance between the Cα atoms of the N- and C-terminal ends indicates if the peptide is in a closed or open conformation. Example conformations are shown in Figure 2A and the overall distribution of all distances over all 3000 ns of all 100 replicas is shown in Figure 2B. It can be seen that the WT and well as the MT spend most of their simulation time in a closed or semi-closed configuration. The WT spends more time in extended configurations than the MT (red tail to the right in Figure 2B) but also lower termini distances are possible in the WT than in the MT (red section on the left in Figure 2B). This indicates that the MT carrying Y/Y at the termini is more stable while the WT carrying G/G at the termini allows spatially for tighter configurations than the MT.

**Figure 2:**
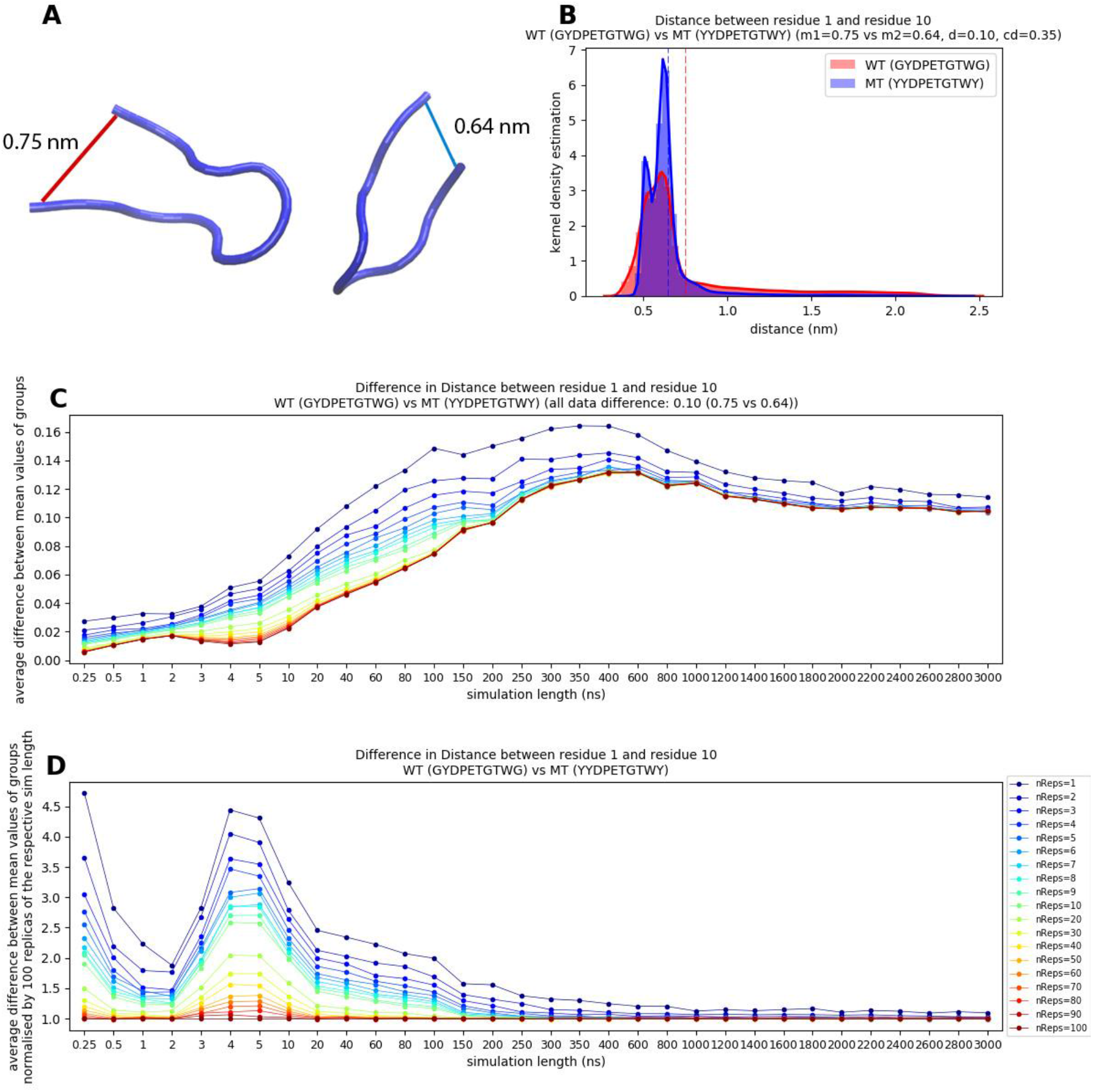
Average difference of the residue1/residue10 Cα distance between wild-type (GYDPETGTWG) and mutant (YYDPETGTWY). (A) 3D representation of example configurations for the average values of subfigure B. The red line shows an example for average value of the red distribution of B while the blue line shows an example for the average value of the blue distribution of B. (B) Distribution of the distances for the WT and MT based on 3000 ns simulations and 100 replicas each. (C) Bootstrapping analysis: Each point represents a unique combination of simulation time and number of replicas. For readability and insight into short simulations the time steps on the x-axis are not equality distributed. (D) Same as (C) but normalised by the value of 100 replicas of the respective simulation time.

The comparison of the average difference between WT and MT at different simulation times and number of replicas based on a bootstrapping analysis is shown in Figure 2C. For the further analysis of the results we assume that 3000 ns and 100 replicas (rightest and lowest dark red point in Figure 2) represent the true difference between WT and MT.

On this basis it can be seen that 100 replicas of very short simulations (0 to 5 ns) strongly underestimate the true difference. Between 5 ns and 600 ns the differences increase until the difference between WT and MT is slightly overestimated from 250 ns onwards. Between 600 ns and 1800 ns the observed difference decreases and becomes stable from about 1800 ns at approximately the assumed true value. For the final 1200 ns the difference remains stable.

For a better visualisation of the replica number influence we have normalised each point of Figure 2C column-wise by the respective difference value of 100 replicas. i.e. 100 replicas represent a difference of 1 and fewer replicas represent a multiple of the 100 replica values. In Figure 2D we show the same data as in Figure 2C but normalised by the value of 100 replicas of each time cut.

It can be seen that few replicas and short simulations show differences that are too large. An increase in replica number and/or simulation time is necessary for reliable results. In simulations longer than 200 ns the use of more than 3-5 replicas does not improve the reliability anymore but an increase in simulation time still improves results until about 1800 ns.

In summary: Simulations of at least 1800 ns using a minimum of 3 replicas should be carried out for reproducible results of the difference between a 10-mer WT and 10-mer MT peptide with respect to the distance between the peptide termini.

### Radius of gyration of WT and MT

The radius of gyration quantifies the compactness of a configuration with respect to its centre of mass. A low radius of gyration indicates a compact structure while a high radius of gyration indicates an extended structure. All atoms are taken into account and each atom is mass weighted.

The radius of gyration of example configurations are schematically illustrated in Figure 3A and the overall distribution of the WT and MT is shown in Figure 3B. In contrast to the peptide termini distance the radius of gyration is larger for the MT than for the WT which is caused by the larger side-chains of the first and last residue in the MT. Even though the radius of gyration measures in principle a similar behaviour as the peptide termini distance also the pattern of the bootstrapping analysis differs considerably (compare Figure 2CD and Figure 3CD). Hundred replicas shorter than 10 ns overestimate the difference in radius of gyration between WT and MT while simulations between 10 ns and 2000 ns underestimate the true difference. Interestingly the use of only two replicas seems to exhibit almost correct differences from 20 ns onwards but this is most likely the result of two errors (too few replicas and too short simulation) cancelling each other out by coincidence.

**Figure 3:**
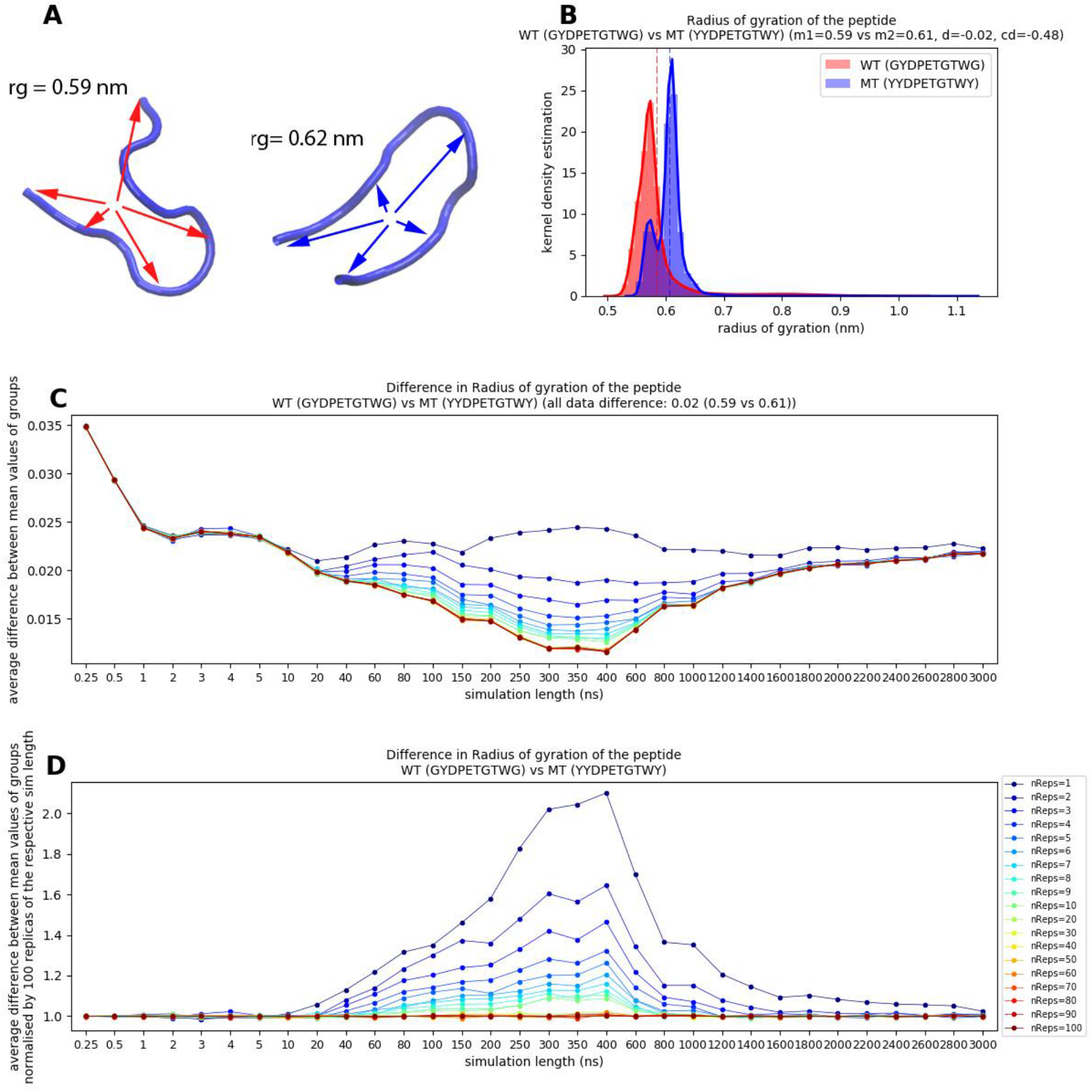
Average difference of the radius of gyration between GYDPETGTWG and YYDPETGTWY. (A) Schematic illustration of the radius of gyration. Arrows indicate distances to the centre of mass. For the actual calculation the distance between the centre of mass and each atom is taken into account. The two configurations represent examples for the mean values of subfigure B. (B) Distribution of the radius of gyration for the WT and MT based on 3000 ns simulations and 100 replicas each. (C) Bootstrapping analysis: Each point represents a unique combination of simulation time and number of replicas. For readability and insight into short simulations the time steps on the x-axis are not equality distributed. (D) Same as (C) but normalised by the value of 100 replicas of the respective simulation time.

Figure 3D shows that the difference between different replicas is large between 10 ns and about 1600 ns. Simulations shorter than 10 ns seem to sample the same local configuration space while simulations longer than 1600 ns provide sufficient sampling that 3 to 5 replicas are sufficient for reproducible differences.

In summary, simulations of more than 2000 ns using a minimum of 3 replicas should be carried out for reproducible results of the difference between a 10-mer WT and 10-mer MT peptide with respect to the radius of gyration.

### Peptide WT and MT: more descriptors

In the supporting material we show the same type of WT/MT difference analysis as in Figure 3 for several other peptide descriptors including the distance between residue 1 and 5 (Figure S 1), distance between residue 6 and 10 (Figure S 2), number of hydrogen bonds within the peptides (Figure S 3), root mean square deviation from the starting structure (Figure S 4), solvent accessible surface area (Figure S 5) and random input (Figure S 6).

The curves of the C-subfigures usually follow a pattern of changing between overestimation and underestimation of the effect or vice versa. This means that short simulations usually start with too high or too low differences and during the course of the simulations this behaviour can change until the correct amount of difference is reached. E.g. Five out of seven descriptors start with an underestimation while purely random data exclusively overestimates the effect (which is an expected finding as the true value is 0). This behaviour together with the recommended simulation time and replicas for different descriptors are summarised in Table 2. Based on this table the general recommendation for additional descriptors with unknown bootstrapping pattern is roughly 3 replicas with a minimum simulation time of 2400 ns. An exception is the SASA (Figure S 5) where the effect size is by far largest and one replica seems sufficient.

**Table 2:** Overview of descriptors for Peptide WT and peptide MT. In the “curve shape” column “u” stands for underestimation, “o” for overestimation, “s” for stable and parenthesis for a weak version of the before mentioned. These estimates are based on the lines represented by many replicas (yellow to red lines). Effect sizes larger than 0.20 are highlighted in bold.

| Descriptor | Curve shape | \|Effect size\| | Simulation time recommended (ns) | Replicas recommended | Figure |
| --- | --- | --- | --- | --- | --- |
| distance(r1/r10) | u-o-s | **0.35** | 1800 | 2-3 | Figure 2 |
| Radius of gyration | o-u-s | **0.48** | 2000 | 2-3 | Figure 3 |
| distance (r1/r5) | u-o-(s) | 0.11 | 2400 | 2-3 | Figure S 1 |
| distance (r6/r10) | o-(u)-s | **0.23** | 800 | 2-3 | Figure S 2 |
| h-bonds within peptide | u-(o)-s | **0.79** | 800 | 1-2 | Figure S 3 |
| RMSD to start | u-o-u-(s) | **0.24** | 2400 | 3-5 | Figure S 4 |
| SASA of the peptide | o-u-s | **1.70** | 1800 | 1 | Figure S 5 |
| random data | o-s | 0.00 | 60 | 1 | Figure S 6 |

The effect size (Table 2) seems to be uncorrelated with the simulation time needed to achieve stable results (r=+0.13 with random data and r=-0.17 without).

### Results of the TCRpMHC systems

The difference between a peptide wild-type and mutant is an interesting setup as excellent coverage of the phase space can be assumed due to the small size of the systems. However, such small peptide systems are rather toy examples as most interesting structures are usually considerably larger than 10 amino acids. We therefore investigated a larger 826 amino acid TCRpMHC system. For this purpose we analysed a total of 200 TCR(pMHC) simulations of 100 ns each (100x wild-type LC13 TCR / FLRGRAYGL peptide / HLA-B*08:01 complex and 100x the same complex carrying the mutant peptide FLRGRAAGL).

### TCRpMHC-WT vs TCRpMHC-7A: CDR3 distance

The first descriptor of the TCRpMHC simulations we investigated was local property: the distance between the CDR3 loop of the TCR α-chain (CDR3α) and CDR3 loop of the TCR β-chain (CDR3β). This is a useful property as the CDR3 loops are centrally located over the potentially immunogenic peptide presented by the MHC (Figure 4A). The spatial proximity allows recognition events taking place within these loops. From the 3D visualisation (Figure 4A; left) it can be seen that the Tyrosine side-chain (green) protrudes deeply between the two CDR loops. In the mutant version (7A) the side-chain is not present (Figure 4A; right).

**Figure 4:**
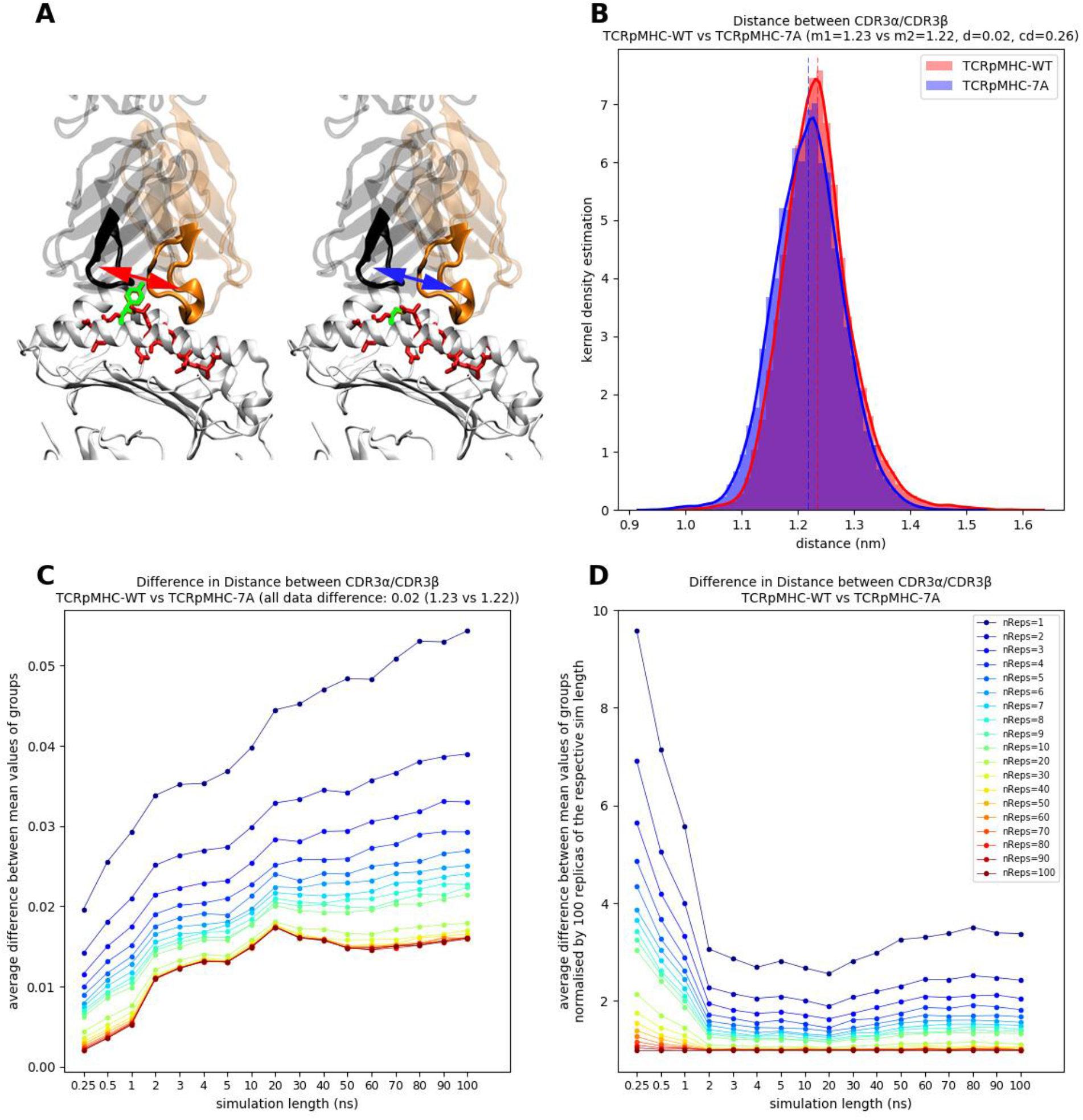
Difference in the CDR3α/CDR3β distance between TCRpMHC wild-type simulations and TCRpMHC mutant (peptide_Y7A_) simulations based on 100 replicas of 100 ns each. (A) Graphical visualisation of the analysed distance. MHC (white), peptide (red), TCR α-chain (transparent orange), CDR3α (orange), TCR β-chain (transparent black), and CDR3β (black), peptide position 7 (green) are shown. Left: TCRpMHC-WT. Right: TCRpMHC-7YA mutant. Red and blue arrows indicate the distance that is analysed. (B) Distribution of the distances during all simulation frames of all replicas. The colour of the arrows in (A) corresponds to the colour of the distributions in (B). (C) Mean difference in CDR3α/CDR3β distance between TCRpMHC wild-type simulations and TCRpMHC mutant simulations based on bootstrapping for different replica numbers and simulation lengths. (D) Same as (C) but normalised column-wise by 100 replicas of the respective simulation length.

This leads to a small difference in the CDR3 distances when comparing the wild-type and mutant simulations (Figure 4B). The difference of 0.02 nm is subtle but Cohen’s d of 0.26 indicates a “small” effect.

While this conclusion seems relatively straight forward from Figure 4B the bootstrapping analysis of Figure 4C shows that the average difference in distance depends strongly on the simulation time and number of replicas. If using only one replica for the WT and 7A then the difference starts at 0.02 nm but quickly increases and does not settle within 100 ns. Using 100 replicas of very short simulations shows hardly any difference but the difference increases quickly up to about 20 ns from which a relatively stable simulation phase with a slight decrease up to 50 ns is reached. This indicates a potential relaxation and readjustment of the binding interface in reaction to the missing Tyrosine residue that then finds its (local) equilibrium again.

Figure 4D shows the same data as Figure 4C but each data point is divided by the value of 100 replicas of this simulation length (i.e. 100 replicas always have a value of 1). It can be seen that very short simulation of up to 2 ns are very dissimilar to each other and one replica on average finds a difference almost 10 times as large as 100 replicas but even in longer simulations few replicas find differences twice as high as 100 replicas.

In summary, simulations of more than 50 ns using a minimum of 10-20 replicas would need to be carried out for reproducible results of the difference between TCRpMHC-WT and the 7A mutant with respect to the distance between CDR3s.

### TCRpMHC-WT vs TCRpMHC-7A: inter TCR chain H-bonds

We next use a more global descriptor of our system: the number of inter TCR chain H-bonds. A change in the number of H-bonds would only be expected if the Y7A mutation destabilises the whole TCR chain arrangement (Figure 5A), however, as the analysis of all 100 replicas for both groups using all 100 ns shows there is no difference in the number of H-bonds between TCRα/ TCRβ and the distributions are almost identical (Figure 5B, mean values of 15.28 and 15.39 H-bonds). Note that this is the same data as we used in our introduction example of Figure 1.

**Figure 5:**
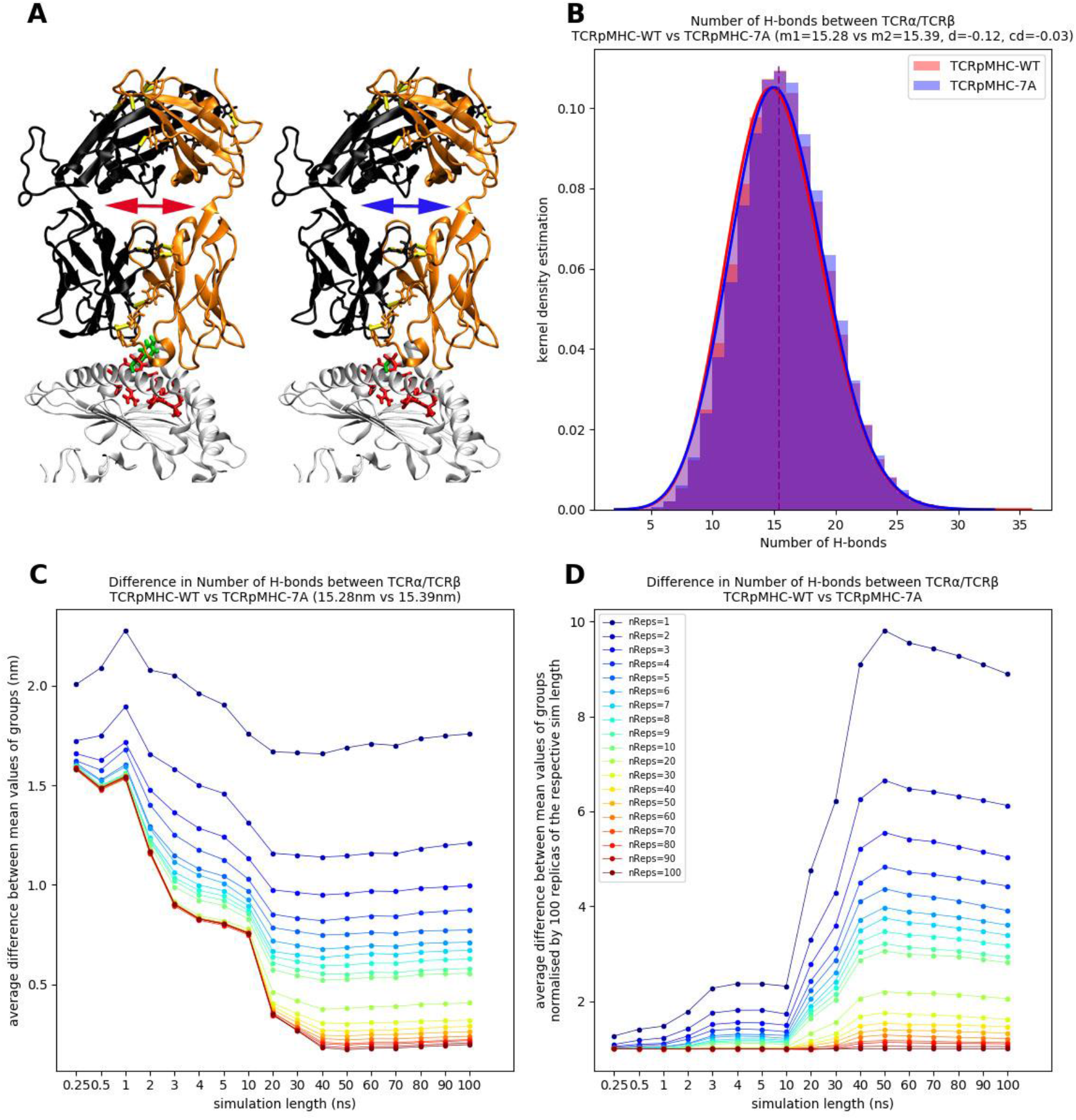
Difference in TCRα/TCRβ number of H-Bonds between TCRpMHC wild-type simulations and TCRpMHC mutant (peptide_Y7A_) simulations based on 100 replicas of 100 ns each. (A) Graphical visualisation of the analysed number of H-bonds. MHC (white), peptide (red), TCR α-chain (orange), TCR β-chain (black), and H-bonds (yellow) are shown. Left: TCRpMHC-WT. Right: TCRpMHC-7A. Red and blue arrows indicate the rough directions of the H-bonds between black and orange. (B) Distribution of the number of H-bonds during all simulation frames of all replicas. The colour of the arrows in (A) corresponds to the colour of the distributions in (B). (C) The mean difference in TCRα/TCRβ number of H-bonds between TCRpMHC wild-type simulations and TCRpMHC mutant simulations based on bootstrapping is shown for different replica numbers and simulation lengths. (D) Same as (C) but normalised column-wise by 100 replicas of the respective simulation length.

Similar to the CDR3 distance the difference in H-bonds converges after about 40 ns to almost 0 if 100 replicas are used (Figure 5C). Again it is important to use at least 10-20 replicas in order to not greatly overestimate the actual effect. Simulations longer than 10 ns show large differences if only few replicas are used (Figure 5D). This peaks in 1vs1 50 ns simulations which overestimate the actual difference by a factor of almost 10. This number is slightly reduced with longer simulations but compared to using more replicas the effect of using longer simulation is small.

In summary, simulations of more than 40 ns using a minimum of 10-20 replicas should be carried out for reproducible results of the difference between TCRpMHC-WT and the 7A mutant with respect to the number of H-bonds between the TCR chains.

### TCRpMHC-WT vs TCRpMHC-7A: more descriptors

In the supporting material we show the same type of TCRpMHC-WT vs TCRpMHC-7A difference analysis for several other descriptors. A summary is given in Table 3. In contrast to the 10-mer peptide system we find a low correlation (r=+0.295) between the effect size and the recommended simulation time to achieve stable results for the TCRpMHC-WT vs TCRpMHC-7A investigation (Table 3). This correlation is mainly based on descriptors with questionable convergence within our simulation time of 10 ns (labelled with “>100” in Table 3) which are mostly the descriptors having larger effect sizes.

**Table 3:** **TCRpMHC-WT and TCRpMHC-7A. In the “curve shape” column “u” stands for underestimation, “o” for overestimation, “s” for stable and parenthesis for a weak version of the before mentioned. These estimates are based on the lines represented by many replicas (yellow to red lines). Effect sizes larger than 0.20 are highlighted in bold.**

| Descriptor | Curve shape | \|Effect size\| | Simulation time recommended (ns) | Replicas recommended | Figure |
| --- | --- | --- | --- | --- | --- |
| CDR3 distance | u-(o)-s | **0.26** | 40 | 10-20 | Figure 4 |
| H-bonds TCR $\alpha/\beta$ | o-s | 0.03 | 50 | 10-20 | Figure 5 |
| H-bonds intra TCR | o-(s) | 0.16 | >100 | 10-20 | Figure S 7 |
| CDR1 distance | (u)-o | **0.36** | >100 | 10-20 | Figure S 8 |
| CDR2 distance | o-u-s | **0.22** | 30 | 20-30 | Figure S 9 |
| RG CDR1α | u-s | **0.32** | 50 | 10-20 | Figure S 10 |
| RG CDR1β | (s) | 0.01 | 5 | 20-30 | Figure S 11 |
| RG CDR2α | o-s | 0.03 | 5 | 20-30 | Figure S 12 |
| RG CDR2β | (s) | 0.06 | 1 | 20-30 | Figure S 13 |
| RG CDR3α | (o)-u-s | 0.16 | 60 | 10-20 | Figure S 14 |
| RG CDR3β | o-u-o-s | 0.07 | 80 | 20-30 | Figure S 15 |
| SASA CDR1α | u-(o)-(s) | **0.21** | >100 | 10-20 | Figure S 16 |
| SASA CDR1β | u-o | 0.09 | >100 | 20-30 | Figure S 17 |
| SASA CDR2α | o-(u)-s | 0.09 | 70 | 20-30 | Figure S 18 |
| SASA CDR2β | u-o | 0.13 | >100 | 20-30 | Figure S 19 |
| SASA CDR3α | o-s | 0.00 | 30 | 20-30 | Figure S 20 |
| SASA CDR3β | u-o-(s) | 0.01 | >100 | 20-30 | Figure S 21 |

## Discussion

In a previous study ^5^ we presented an analysis of intra-replica differences i.e. if the same simulation is performed multiple times: how much simulation time and how many replicas and are needed to obtain on average a difference close to zero between random subgroups of identical replicas. In this previous study we found that about 10 replicas are a good trade-off between reliability of the results and computational time invested. In the current study we go one step further and analyse the same challenge but for inter-simulation differences i.e. two different systems are simulated multiple times but in this case the two groups of simulations are expected to differ from each other and the sought difference is not always zero as for intra-simulation replicas.

Our study quantifies to our knowledge for the first time the actual extent of the variably in inter-simulation differences. This allows us to assess the results of other studies in terms of accuracy and reliability of the reported effect(s). Our study suggests that future comparison studies should consider the use of replicas.

Our results are applicable for standard molecular simulations but there might be a possibility that our results are not applicable to all types of advanced simulation techniques such as replica-exchange simulations, metadynamics, gaussian accelerated molecular dynamics or similar. However we have previously found indications that also advanced sampling techniques that use e.g. hierarchical natural move Monte Carlo suffer from similar problems of non-reproducibility as standard simulations (Figure 5 of ^28^).

In the current study we present analysis of two different systems. The first system was the comparison of two 10 amino acid peptides. This system is small enough that our total simulation time (3000 ns with 100 replicas for the WT and MT) covers the known folding time of 600 ns ^29^ around 500 times. In this case we can assume that we have achieved complete sampling and good statistics about all possible configurations. In such cases few replicas (n∼3) but very long simulations (>2000 ns) seem to be the best choice to determine the correct difference between the WT and MT peptide.

The second system we analysed was an 826 amino acids TCRpMHC complex. For systems of this size a complete sampling of the free energy landscape cannot be expected as current simulation times are not even close to the natural folding time of larger proteins ^30^. Simulations are rather expected to explore one or more local free energy basins around the folded X-ray structure state. Although this is sub-optimal from a statistical point of view this is the usual situation in a molecular simulation project as usual structures of interest are considerably larger than the 10 amino acids and simulation time is always finite. In our larger 826 amino acids TCRpMHC system with its restricted sampling time the picture changes considerably. In general between 10 and 40 replicas are needed in order to obtain reproducible results. This suggests that more replicas are required when testing for inter-simulation differences compared to intra-simulation differences. This is potentially because different replicas explore different local energy basins and this bias is not cancelling out between mutant and wild-type but rather adding up. This yields to a recommendation of a minimum of 10 to 20 replicas.

So do these false-positively found differences between mutant and wild-type actually matter in terms of scale? As an example we take the number of H-bonds between the two TCR chains (Figure 5). This is the same example that was already discussed in the introduction (Figure 1). In the introduction the most extreme case of false-positive reporting was discussed in which the wild-type simulation had 19 H-bonds while the mutant had only 12 H-bonds (the true difference was almost 0). This demonstrates that, single extreme outliers can happen but the more interesting question is the average mistake. In Figure 5 it can be seen that on average the results of single simulations would be almost 2 H bonds off. Even with 3 or 4 replicas each the results are about 1 H-bond off. If we want to bring the mistake down to half an H-bond 10 replicas are sufficient while if a quarter of an H-bond is our limit about 20 replicas are needed. In the end it is a trade-off of between computational time invested and accuracy/reproducibility of the results.

## Conclusion

In general more replicas and simulation time improve the reliability of results but the actual number of replicas and time needed depends on the system and descriptor under investigation. If the simulation length covers the experimental folding time multiple times (which is rarely the case in practice) using as few as 3 replicas can make sense. However, for the vast majority of studies the simulation time is much shorter than the experimental folding time and thus results based on fewer than 10-20 replicas per comparison group will likely exhibit false-positive findings for the majority of molecule comparison studies.

## Data and Software Availability

All data are available from: https://figshare.com/articles/dataset/CLN_1UAO_GYDPETGTWG_wildtype/19793764, https://figshare.com/articles/dataset/CLN025_2RVD_3000ns_mutant/19794073 and https://figshare.com/articles/LC13_tar_gz/8067746.

## Supporting information

**Figure S 1:**
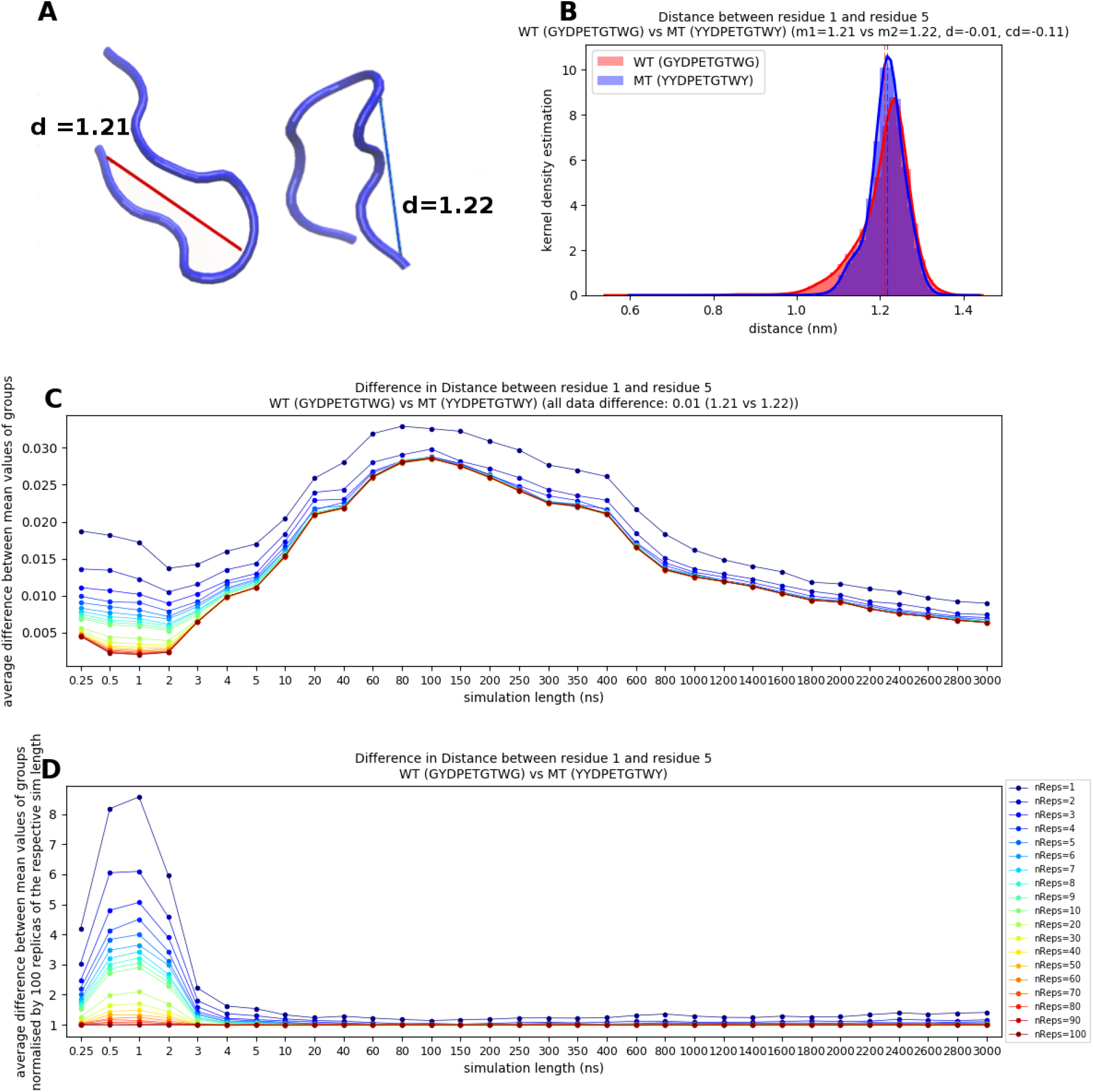
Average difference of the residue1/residue5 Cα distance between wild-type (GYDPETGTWG) and mutant (YYDPETGTWY). (A) 3D representation of example configurations for the average values of subfigure B. (B) Distribution of the distances for the WT and MT based on 3000 ns simulations and 100 replicas each. (C) Bootstrapping analysis: Each point represents a unique combination of simulation time and number of replicas. For readability and insight into short simulations the time steps on the x-axis are not equality distributed. (D) Same as (C) but normalised by the value of 100 replicas of the respective simulation time.

**Figure S 2:**
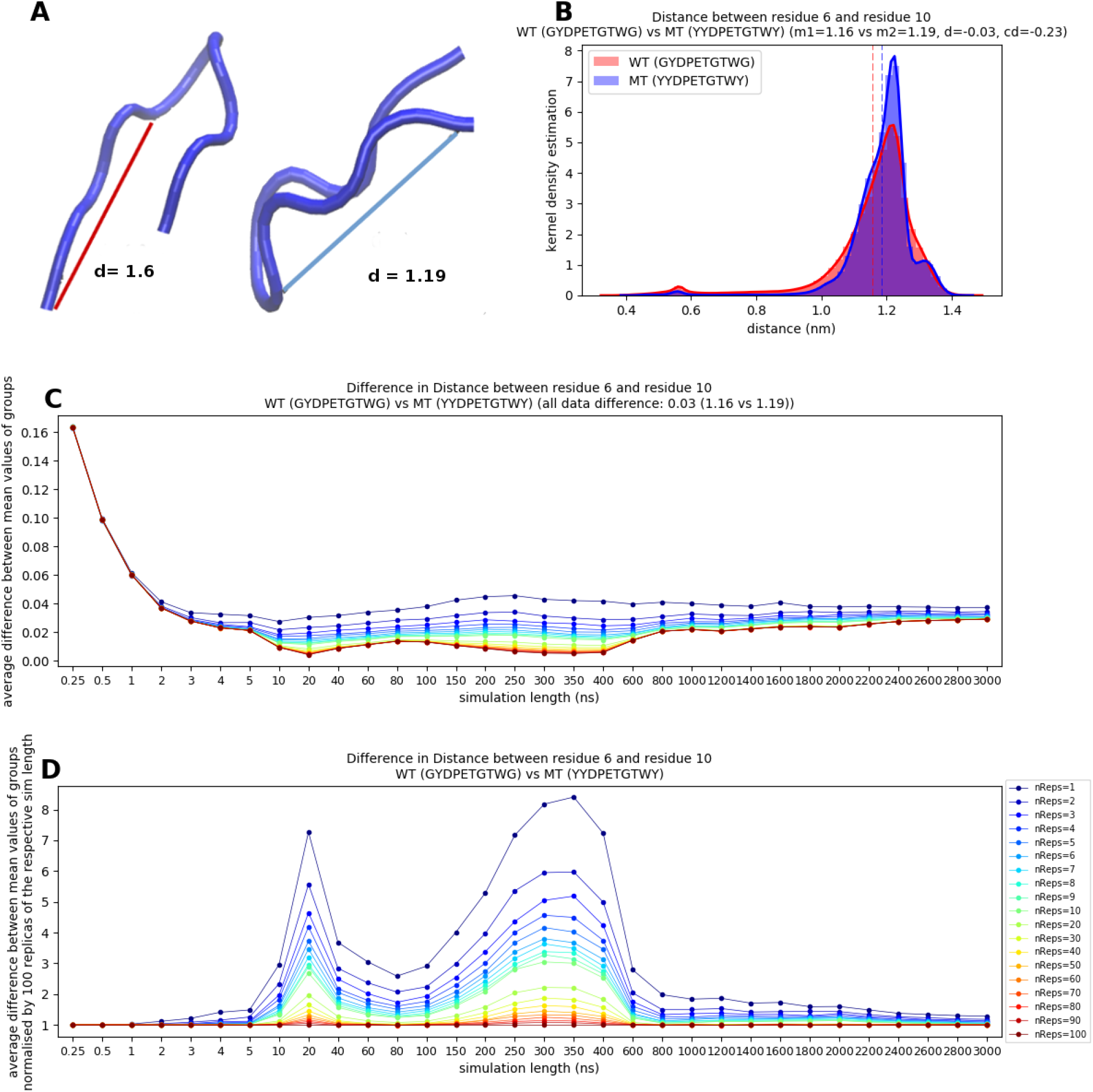
Average difference of the residue6/residue10 Cα distance between wild-type (GYDPETGTWG) and mutant (YYDPETGTWY). (A) 3D representation of example configurations for the average values of subfigure B. (B) Distribution of the distances for the WT and MT based on 3000 ns simulations and 100 replicas each. (C) Bootstrapping analysis: Each point represents a unique combination of simulation time and number of replicas. For readability and insight into short simulations the time steps on the x-axis are not equality distributed. (D) Same as (C) but normalised by the value of 100 replicas of the respective simulation time.

**Figure S 3:**
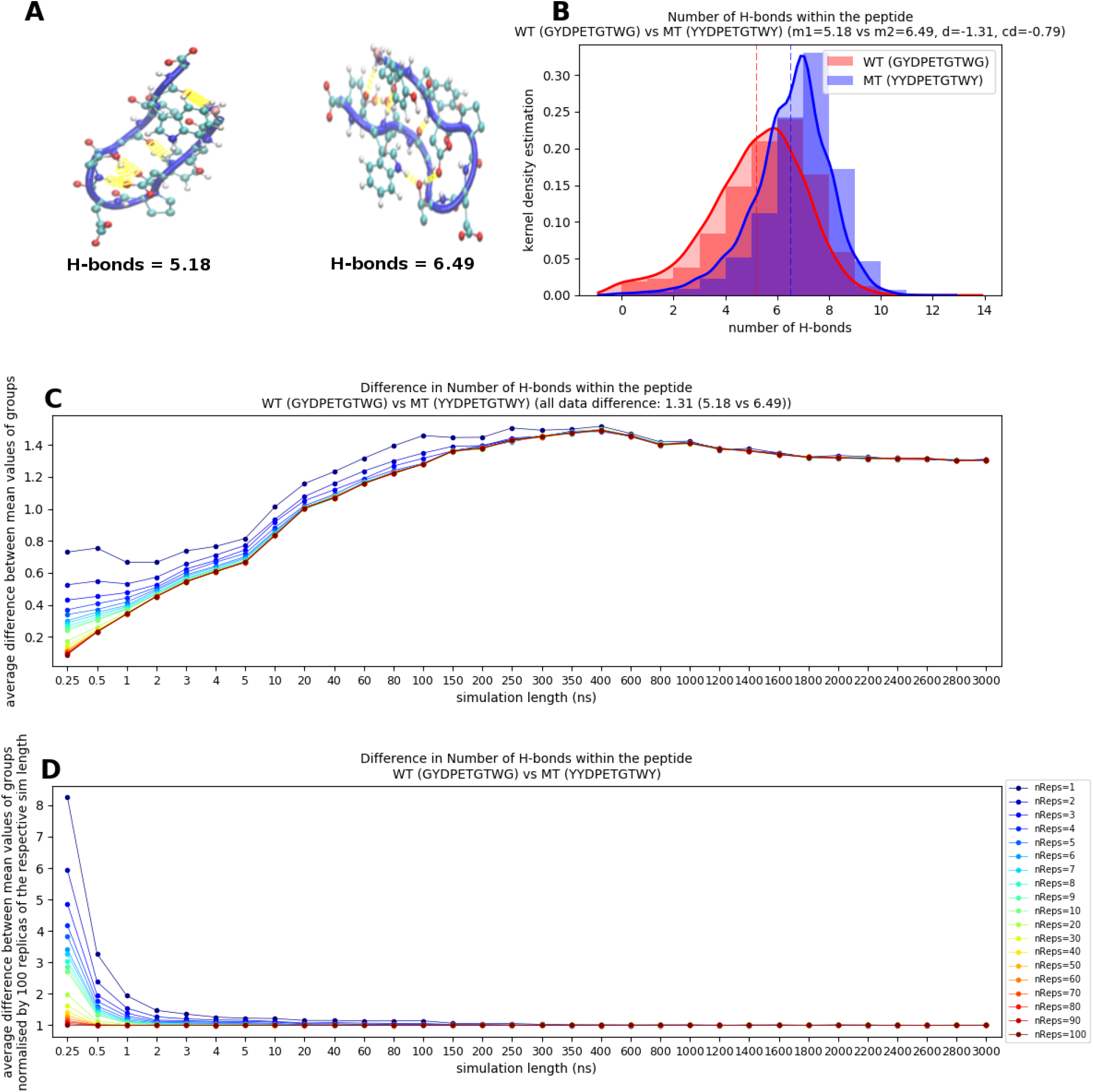
Average difference of the number of hydrogen bonds between wild-type (GYDPETGTWG) and mutant (YYDPETGTWY). (A) 3D representation of example configurations for the average values of subfigure B. The hydrogen bonds are shown in yellow. (B) Distribution of the H-bonds for the WT and MT based on 3000 ns simulations and 100 replicas each. (C) Bootstrapping analysis: Each point represents a unique combination of simulation time and number of replicas. For readability and insight into short simulations the time steps on the x-axis are not equality distributed. (D) Same as (C) but normalised by the value of 100 replicas of the respective simulation time.

**Figure S 4:**
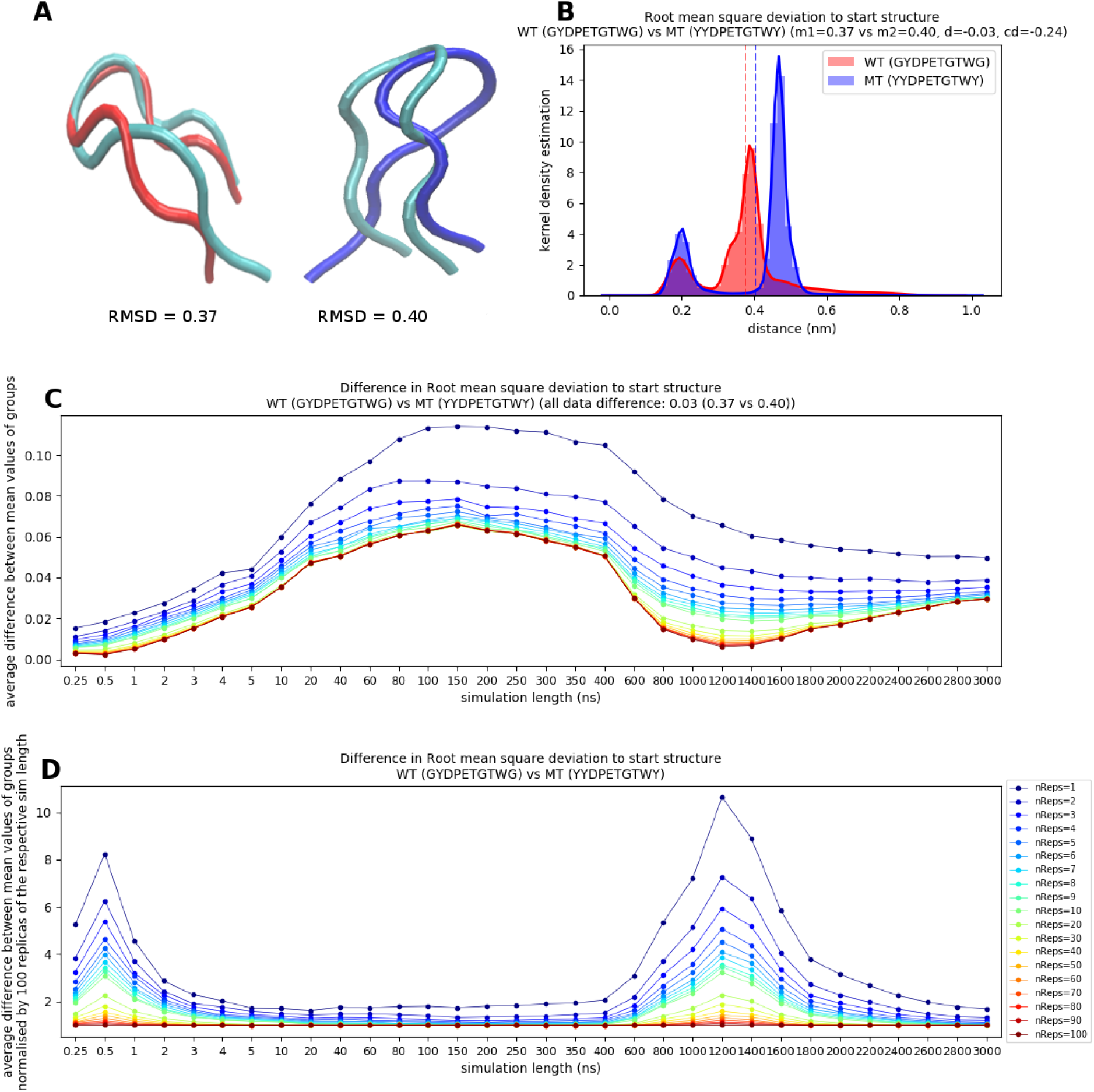
Average difference of the root mean square deviation between wild-type (GYDPETGTWG) and mutant (YYDPETGTWY). (A) 3D representation of example configurations for the average values of subfigure B. The x-ray configuration is shown in cyan. (B) Distribution of the RMSD for the WT and MT based on 3000 ns simulations and 100 replicas each. (C) Bootstrapping analysis: Each point represents a unique combination of simulation time and number of replicas. For readability and insight into short simulations the time steps on the x-axis are not equality distributed. (D) Same as (C) but normalised by the value of 100 replicas of the respective simulation time.

**Figure S 5:**
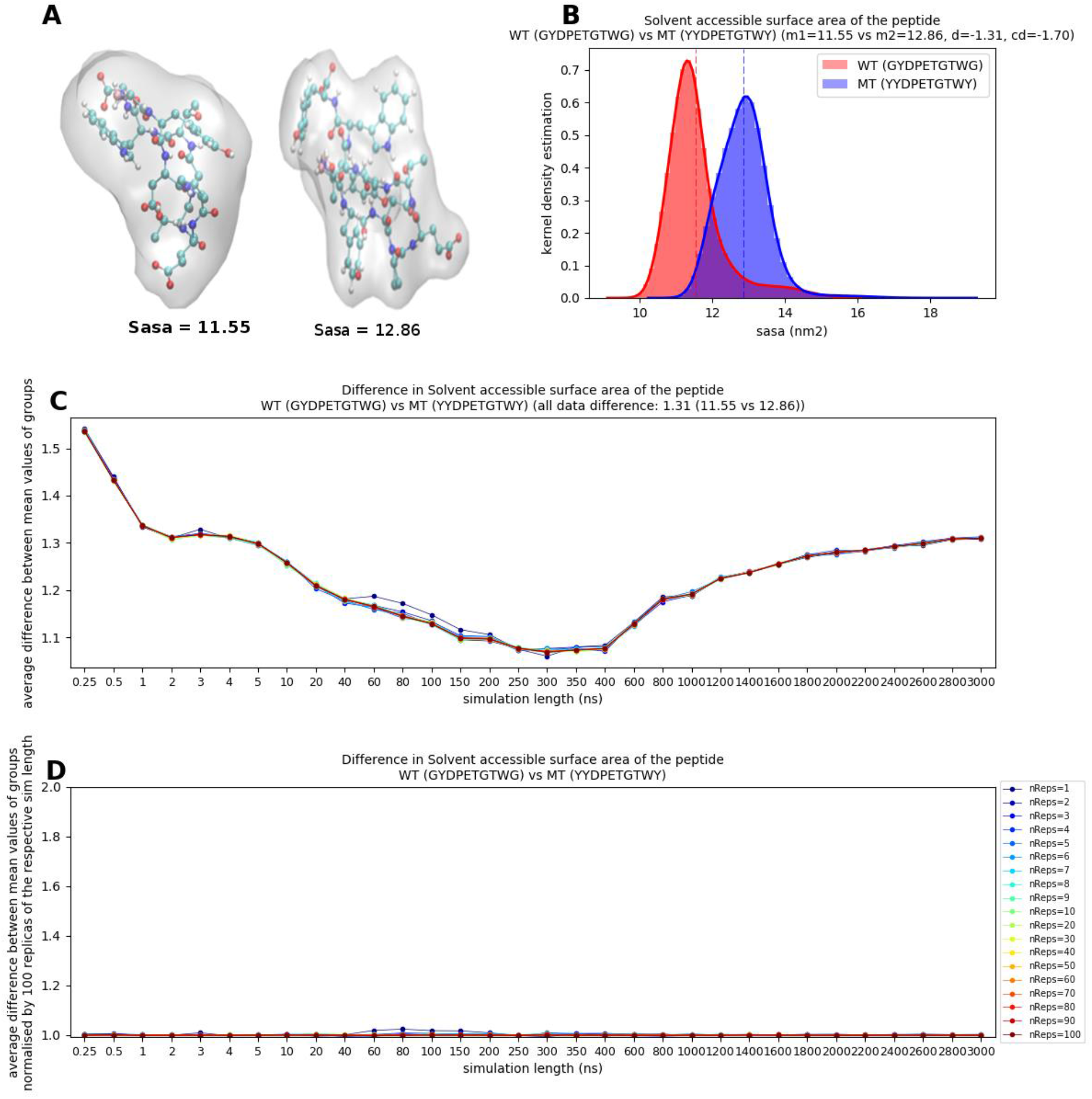
Average difference of the solvent accessible surface area between wild-type (GYDPETGTWG) and mutant (YYDPETGTWY). (A) 3D representation of example configurations for the average values of subfigure B. The SASA is shown as grey transparent surface. (B) Distribution of the SASA for the WT and MT based on 3000 ns simulations and 100 replicas each. (C) Bootstrapping analysis: Each point represents a unique combination of simulation time and number of replicas. For readability and insight into short simulations the time steps on the x-axis are not equality distributed. (D) Same as (C) but normalised by the value of 100 replicas of the respective simulation time.

**Figure S 6:**
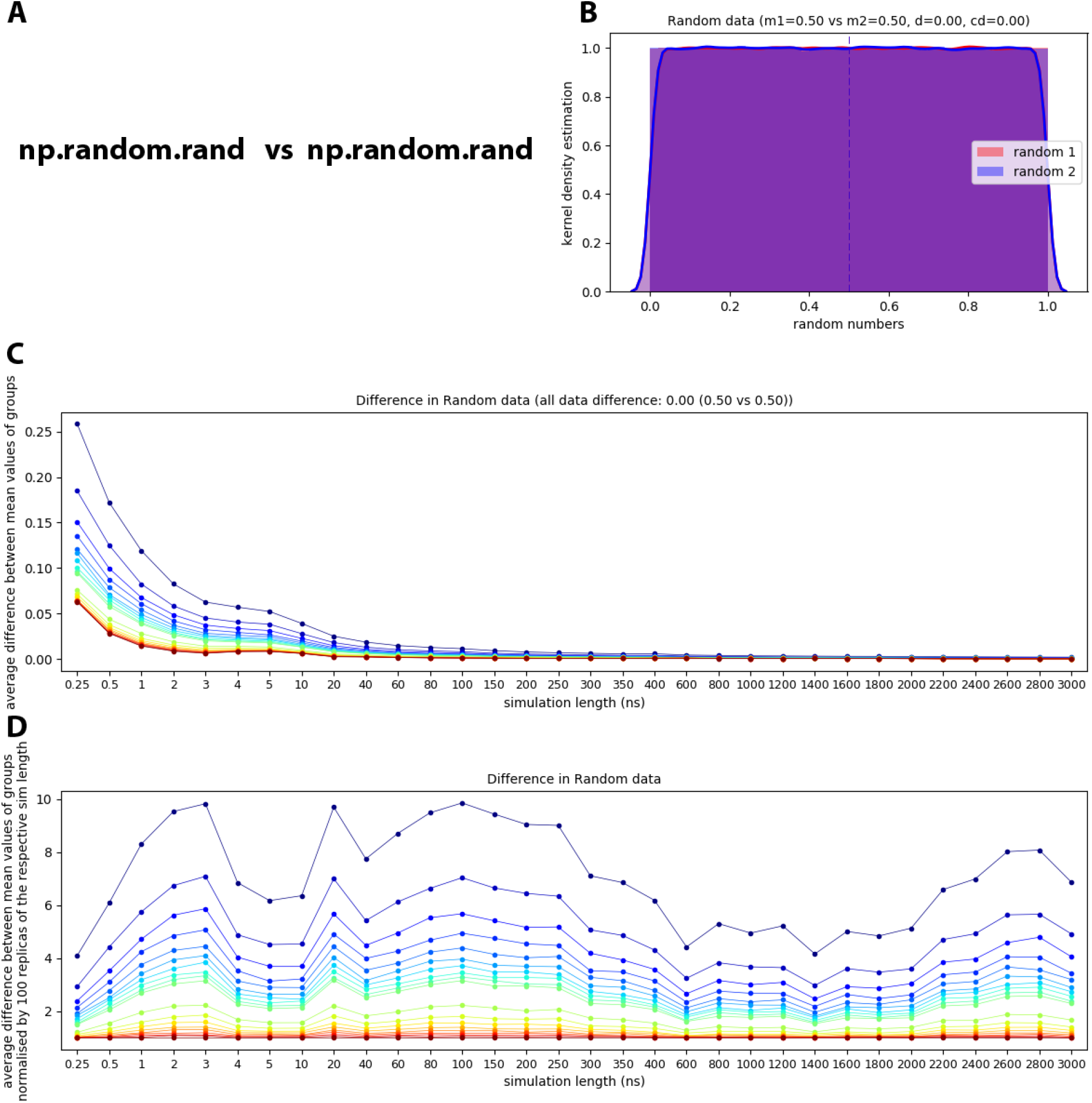
Average difference of the residue1/residue5 Cα distance between wild-type (GYDPETGTWG) and mutant (YYDPETGTWY). (A) These data was generated using the random function of numpy. (B) Distribution of the data. (C) Bootstrapping analysis: Each point represents a unique combination of simulation time and number of replicas. For readability and insight into short simulations the time steps on the x-axis are not equality distributed. (D) Same as (C) but normalised by the value of 100 replicas of the respective simulation time.

**Figure S 7:**
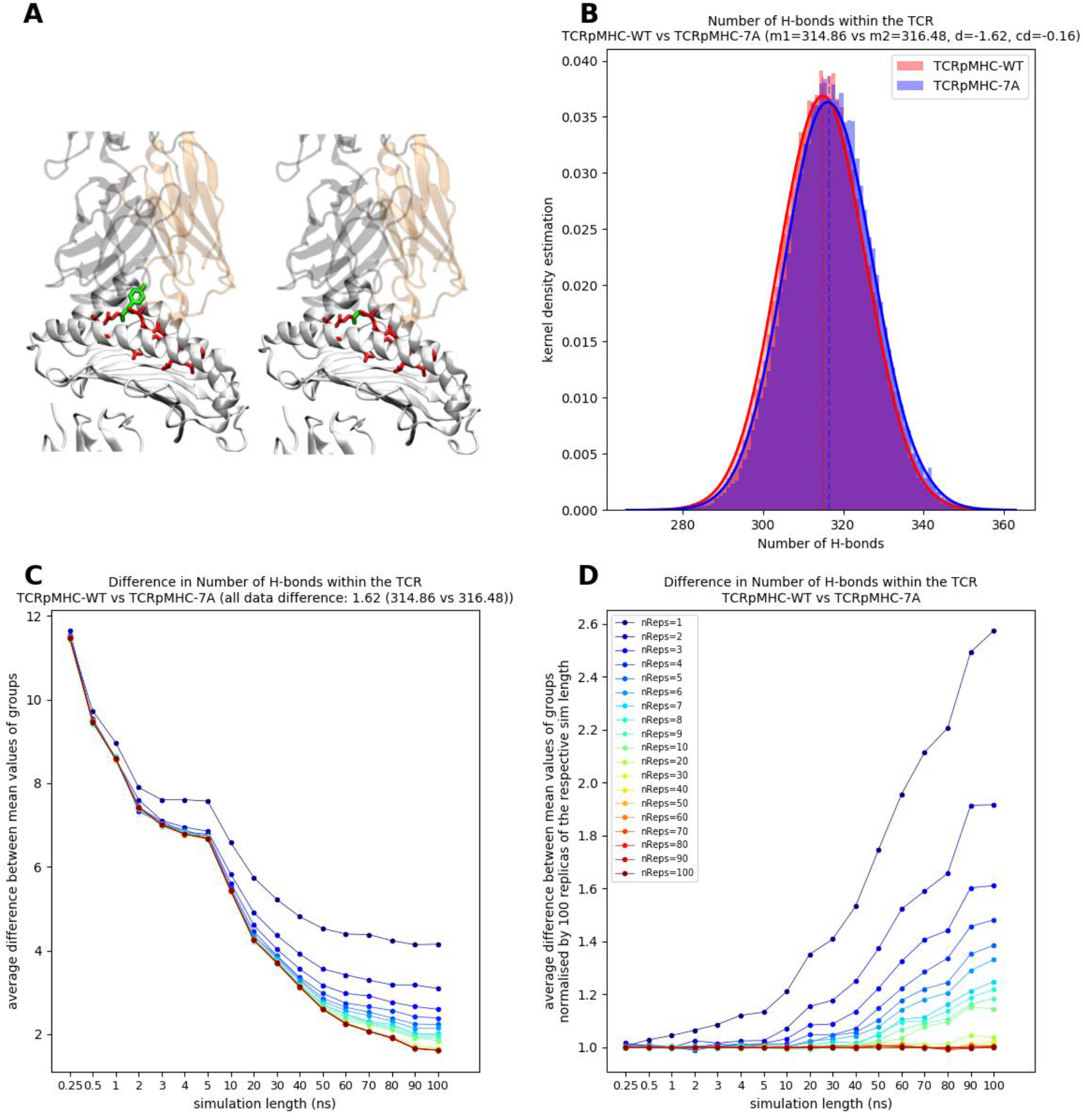
Same as Figure 4 but for intra-TCR H-bonds.

**Figure S 8:**
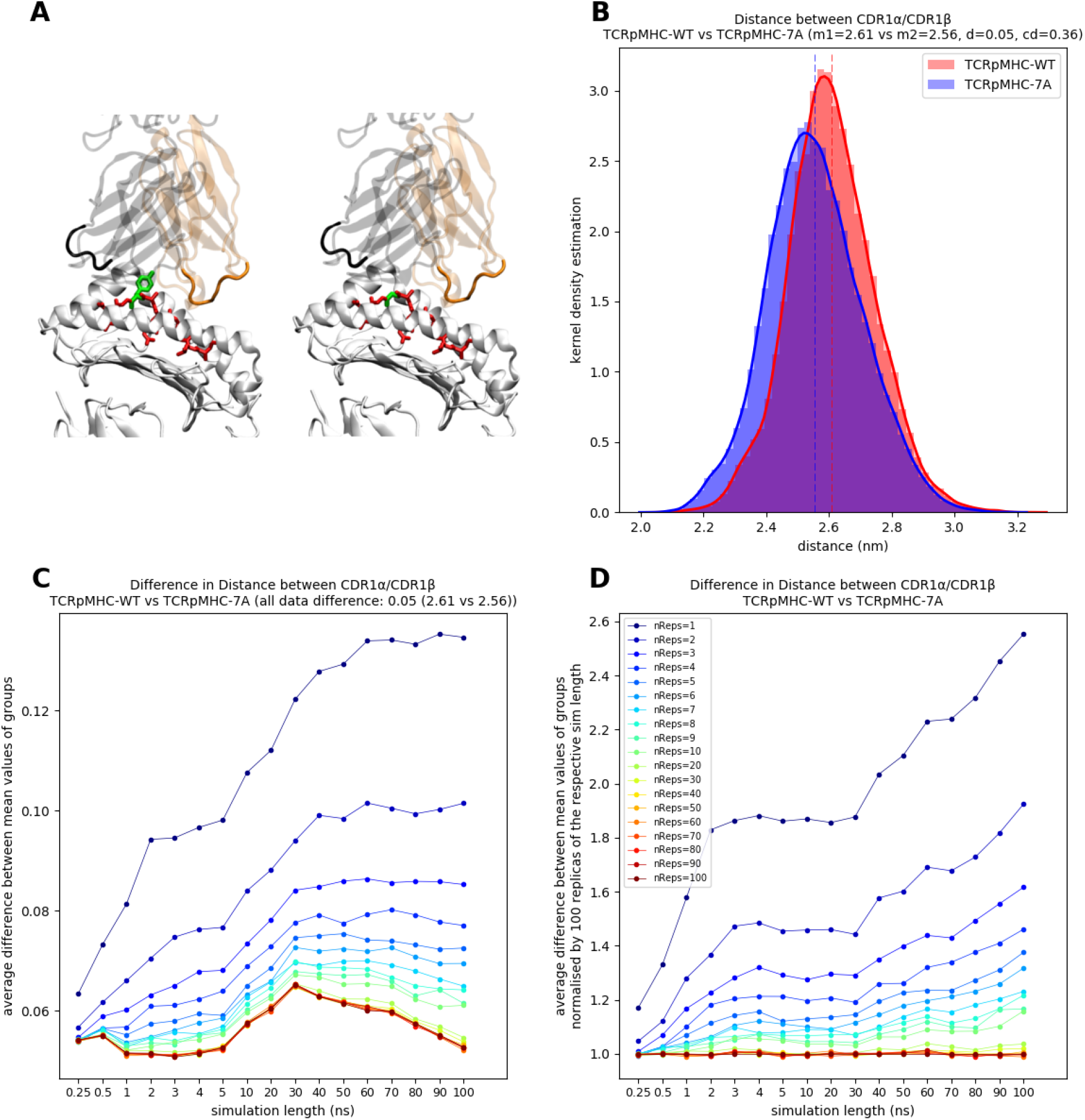
Same as Figure 4 but for the distance between CDR1α and CDR1β.

**Figure S 9:**
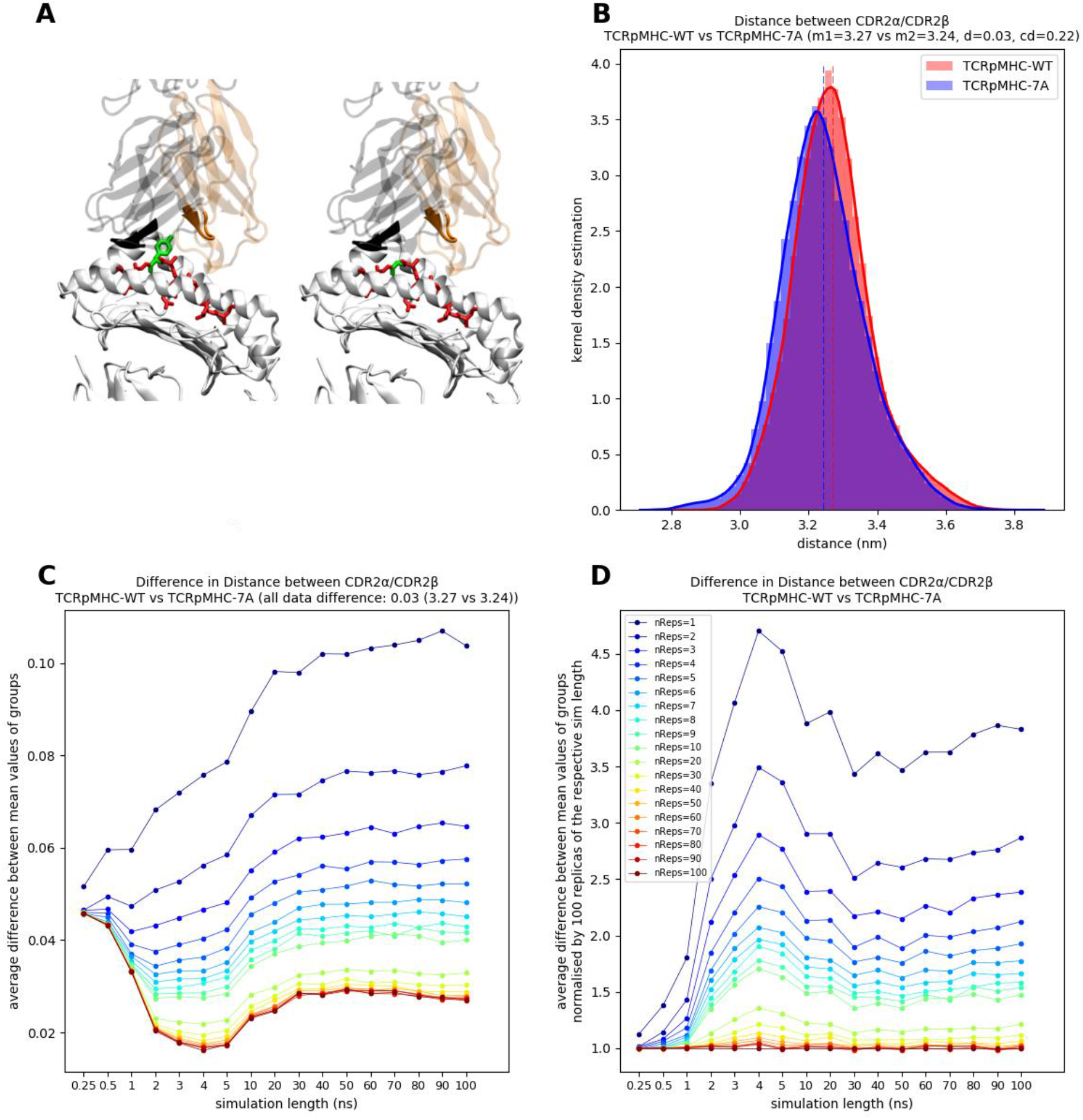
Same as Figure 4 but for the distance between CDR2α and CDR2β.

**Figure S 10:**
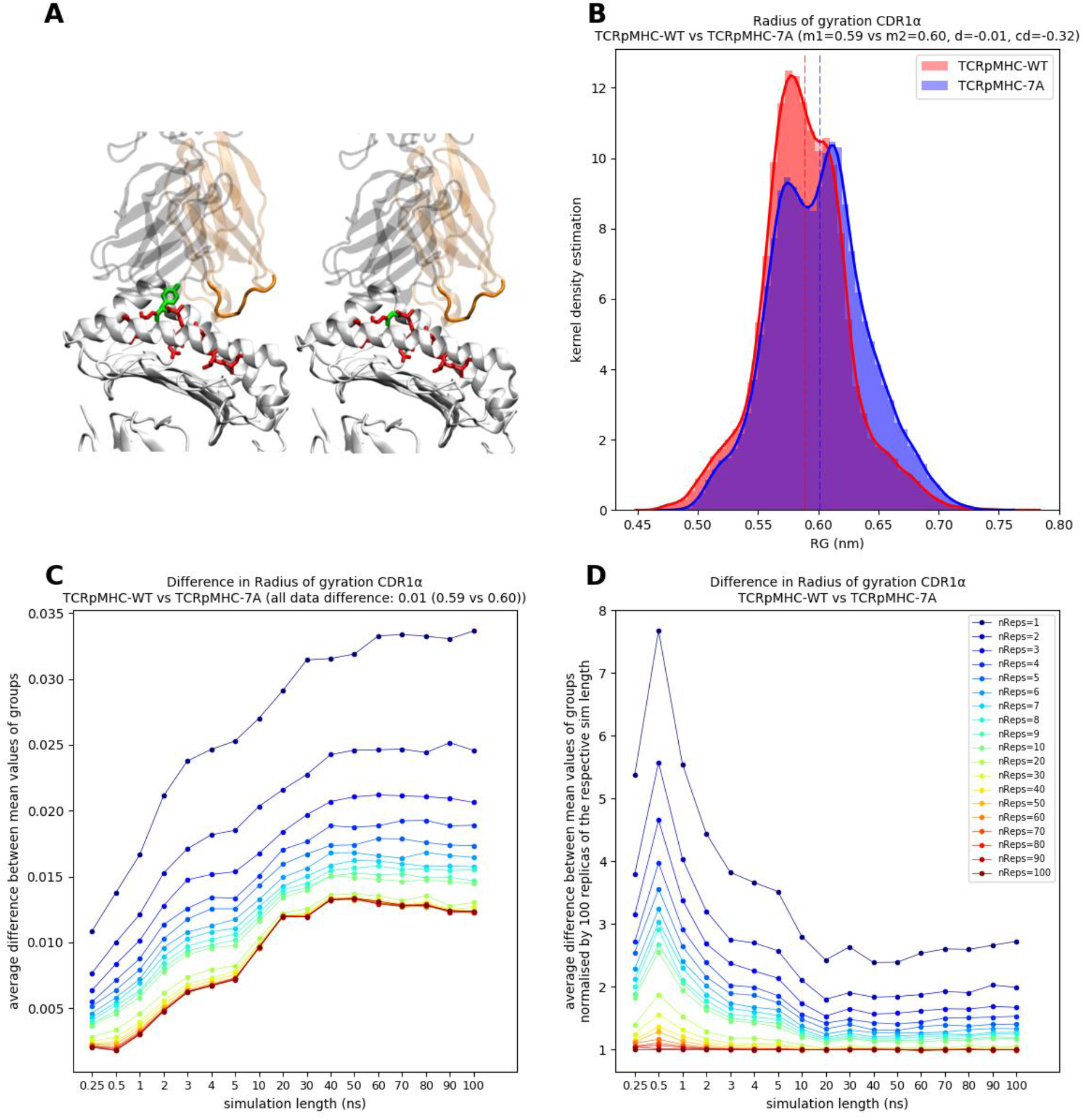
Same as Figure 4 but for the radius of gyration of CDR1α.

**Figure S 11:**
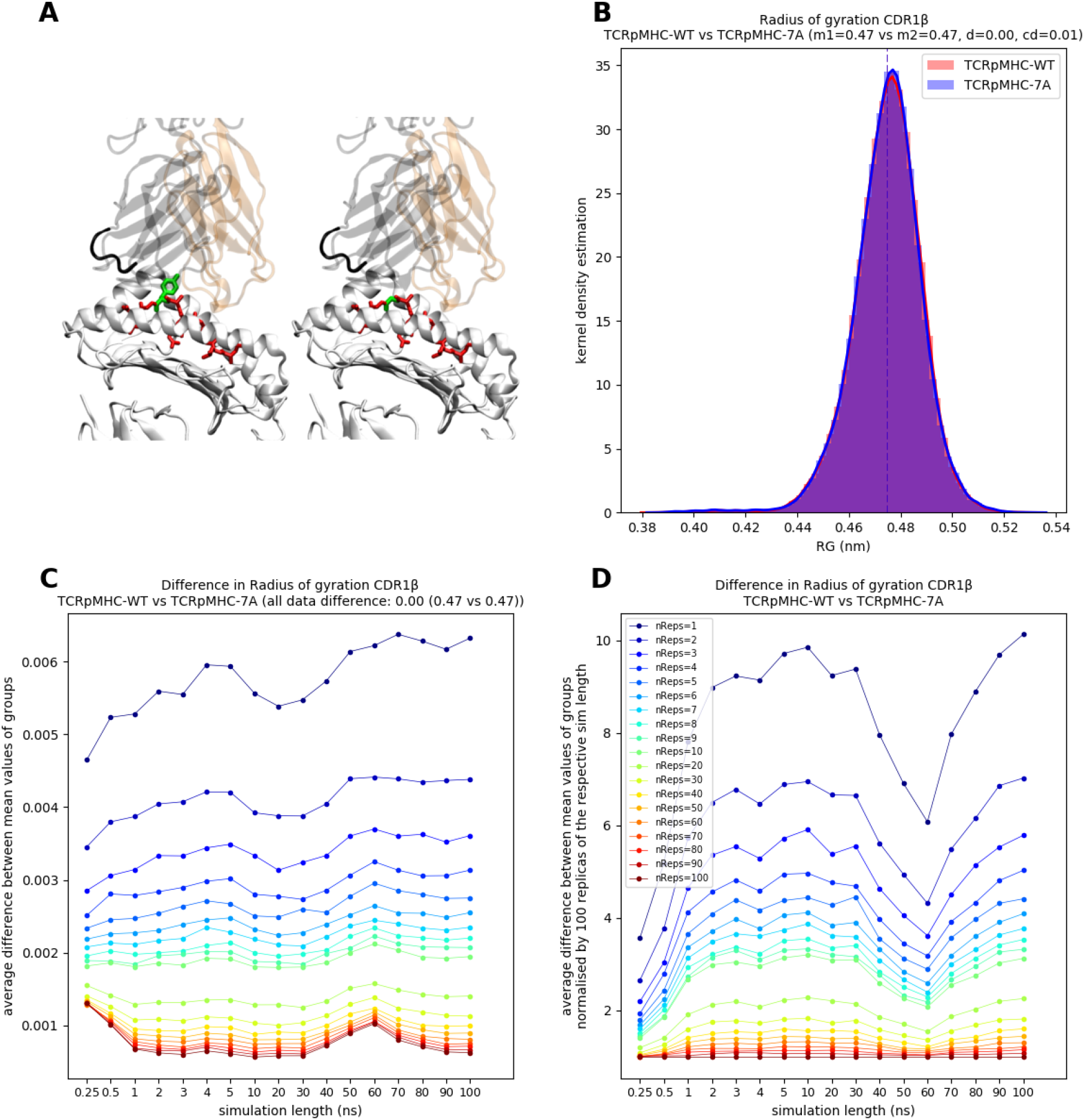
Same as Figure 4 but for the radius of gyration of CDR1β.

**Figure S 12:**
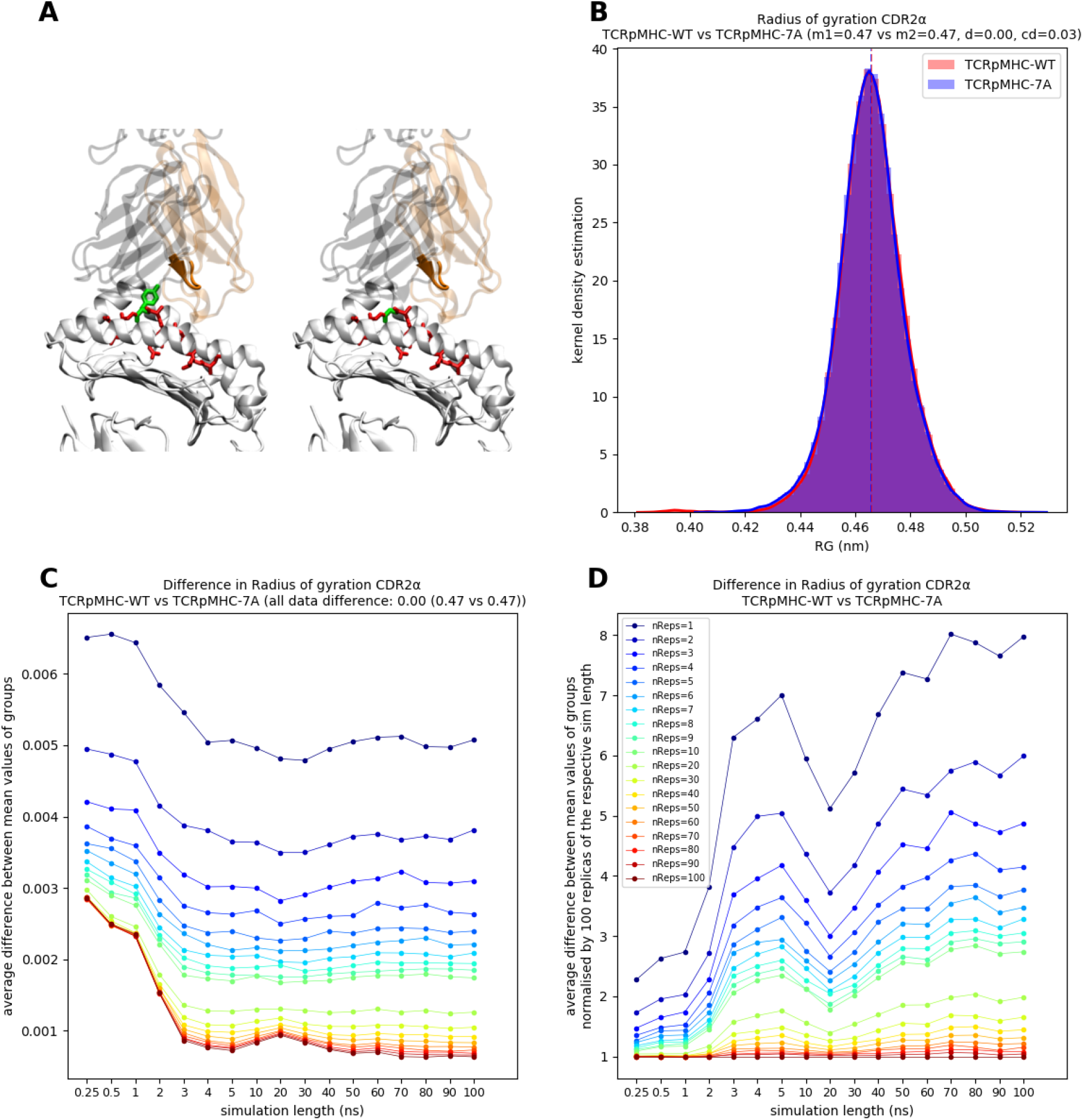
Same as Figure 4 but for the radius of gyration of CDR2α.

**Figure S 13:**
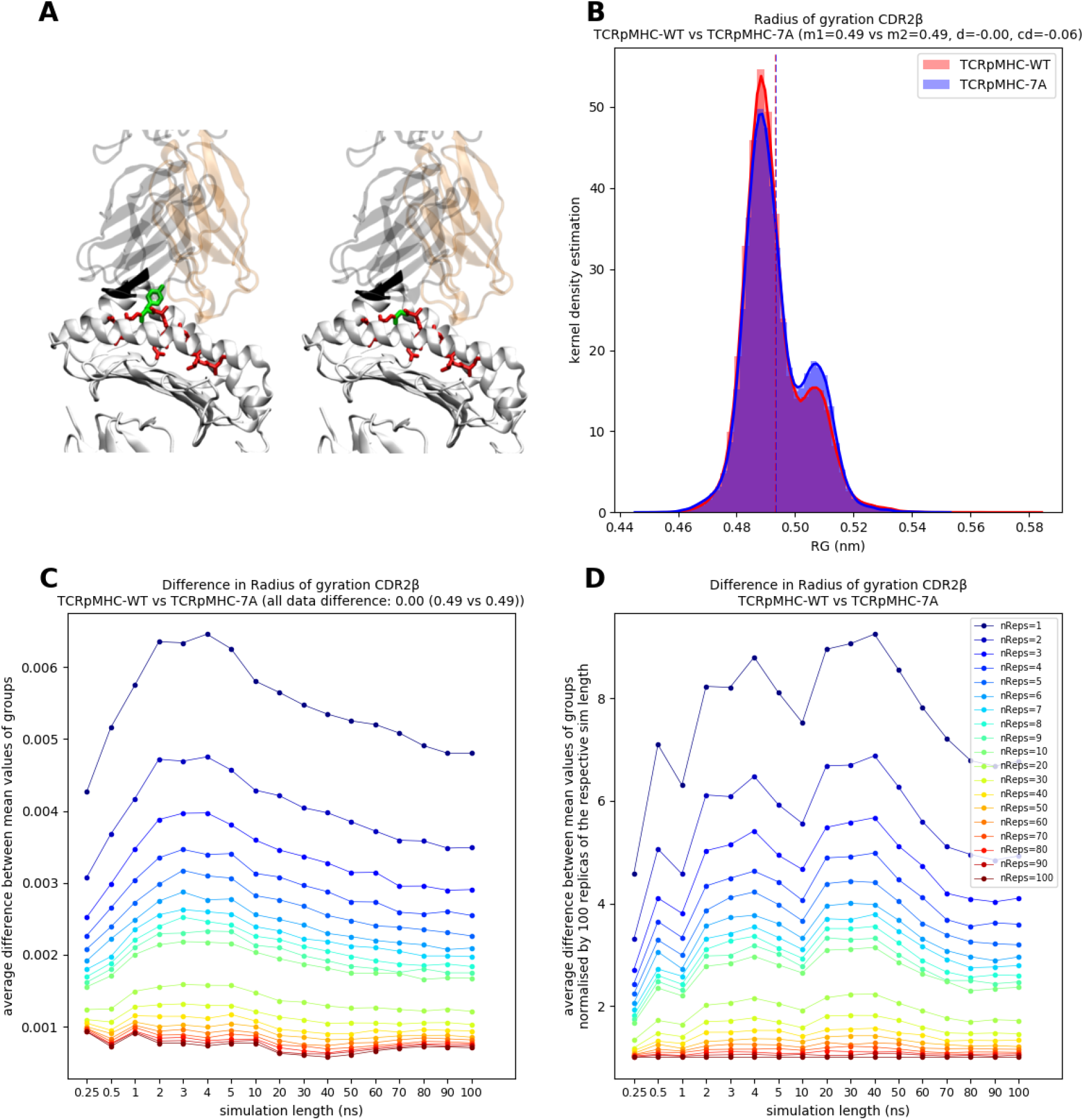
Same as Figure 4 but for the radius of gyration of CDR2β.

**Figure S 14:**
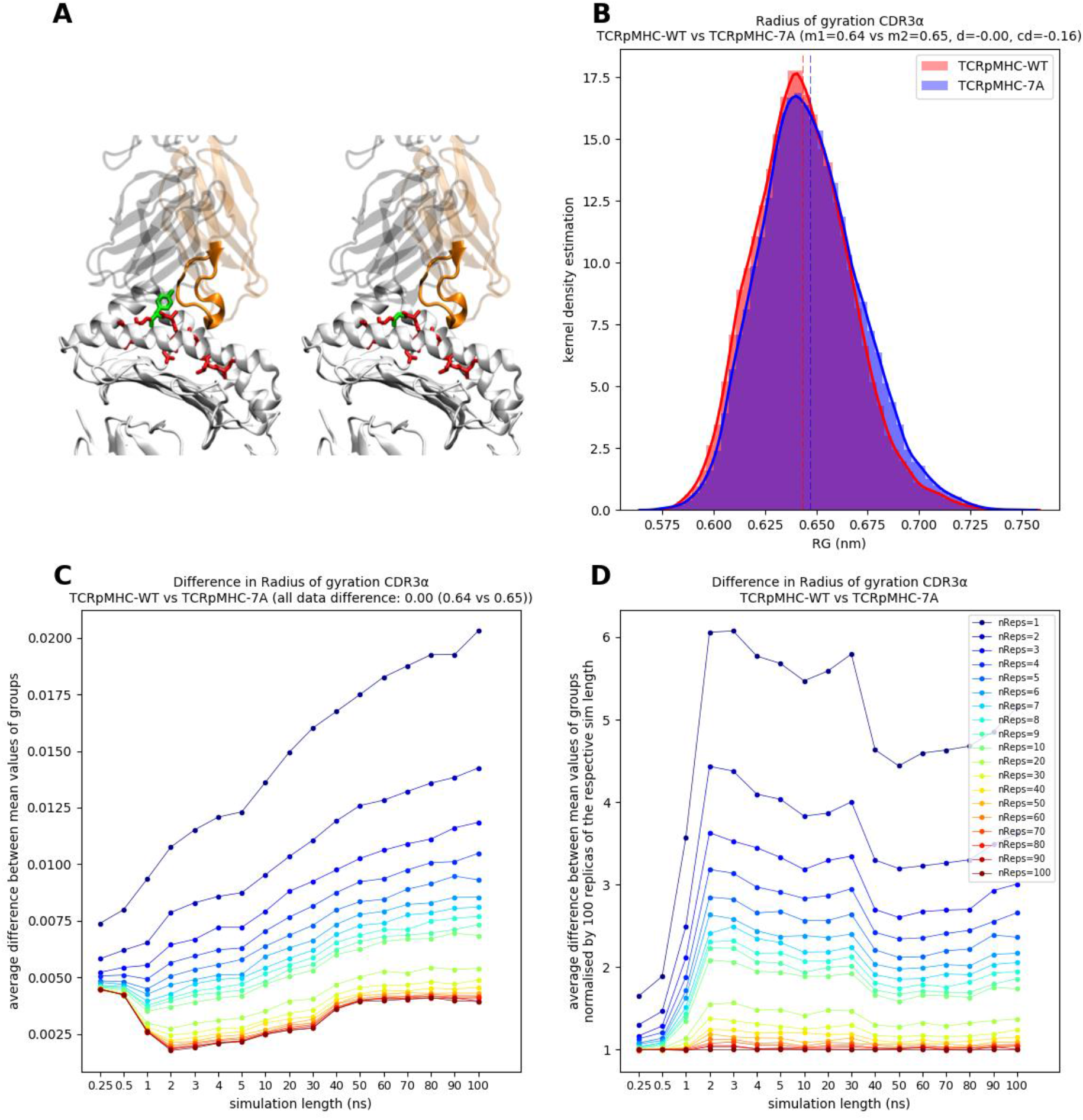
Same as Figure 4 but for the radius of gyration of CDR3α.

**Figure S 15:**
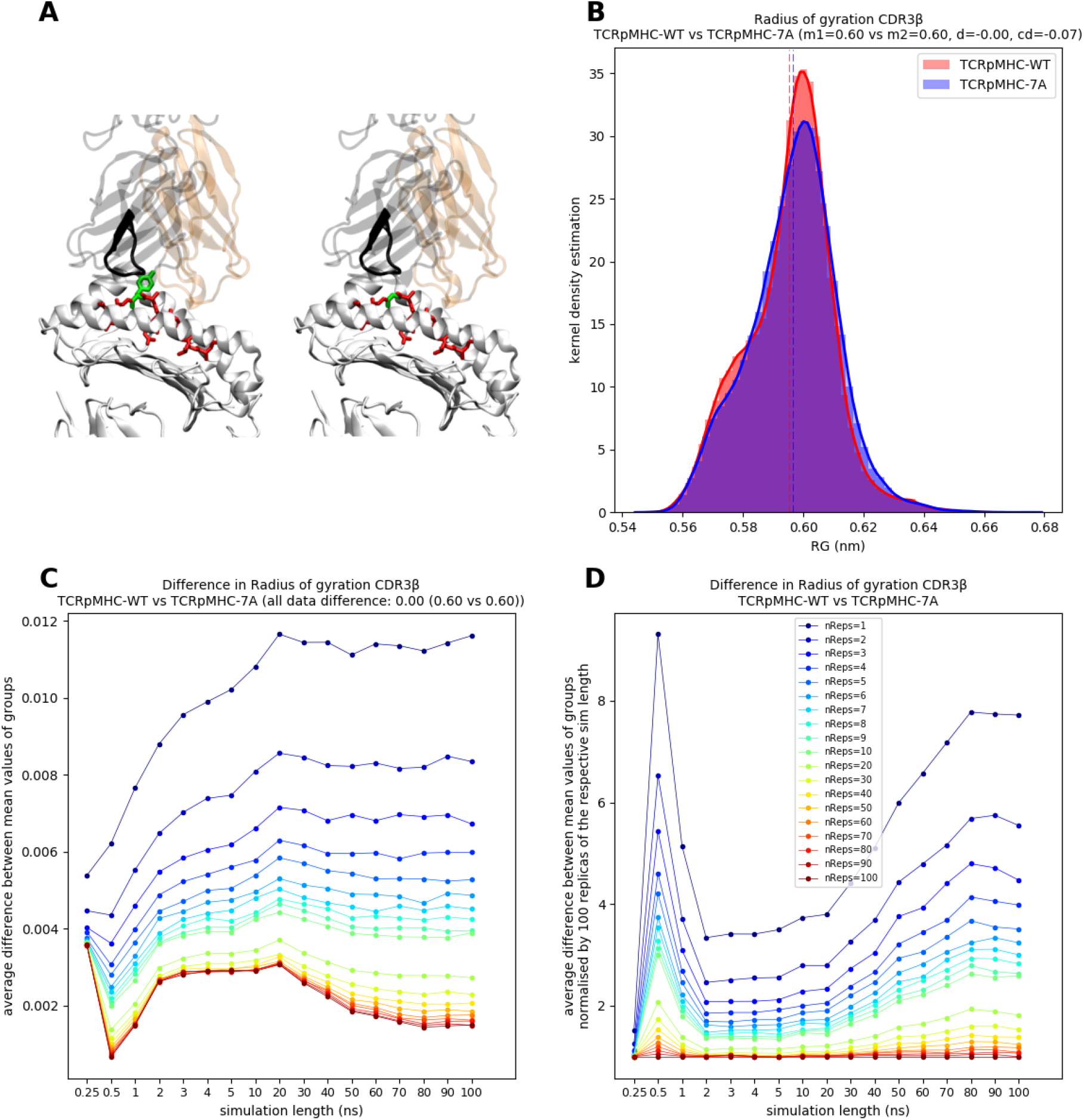
Same as Figure 4 but for the radius of gyration of CDR3β.

**Figure S 16:**
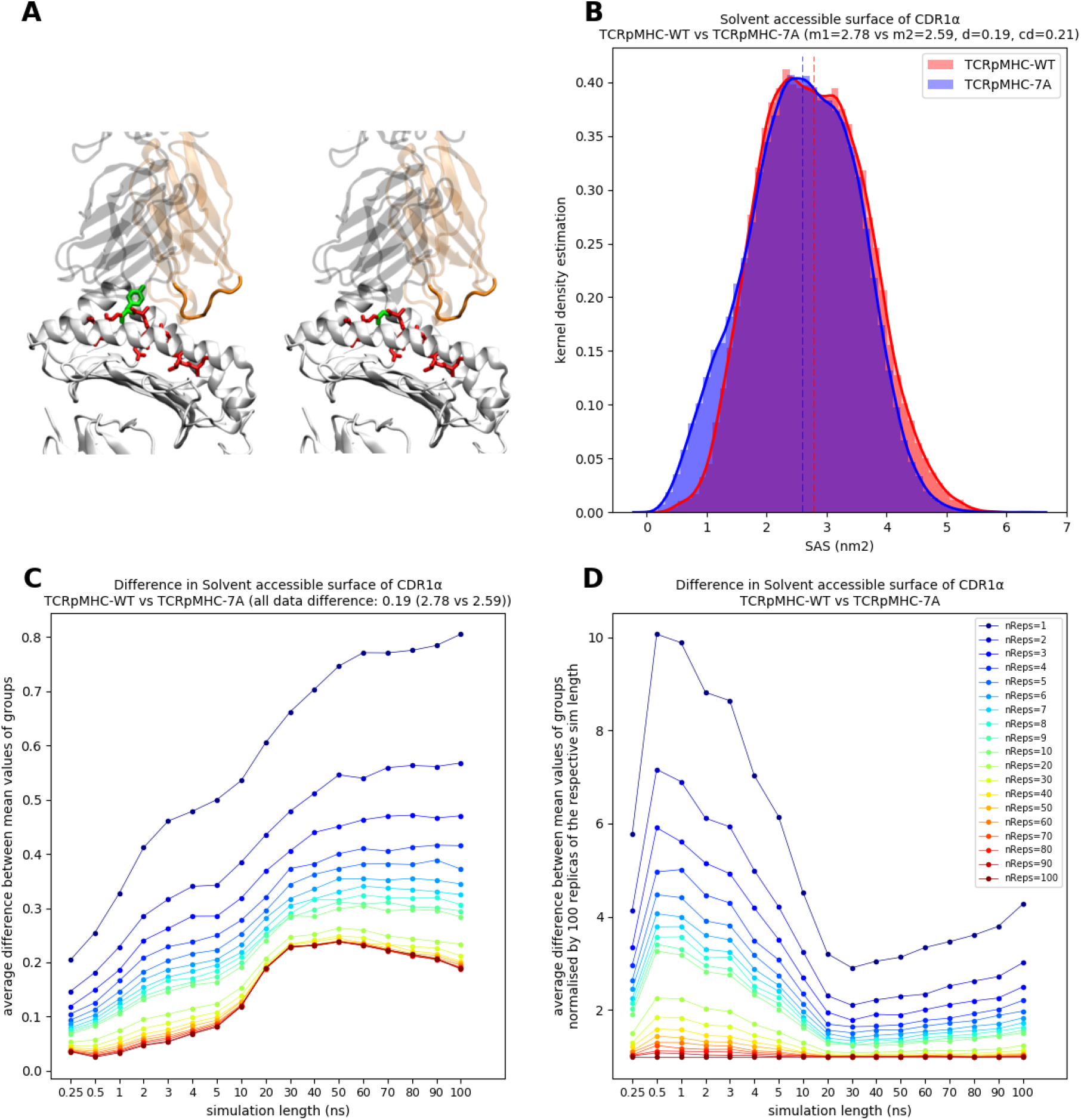
Same as Figure 4 but for the SASA of CDR1α.

**Figure S 17:**
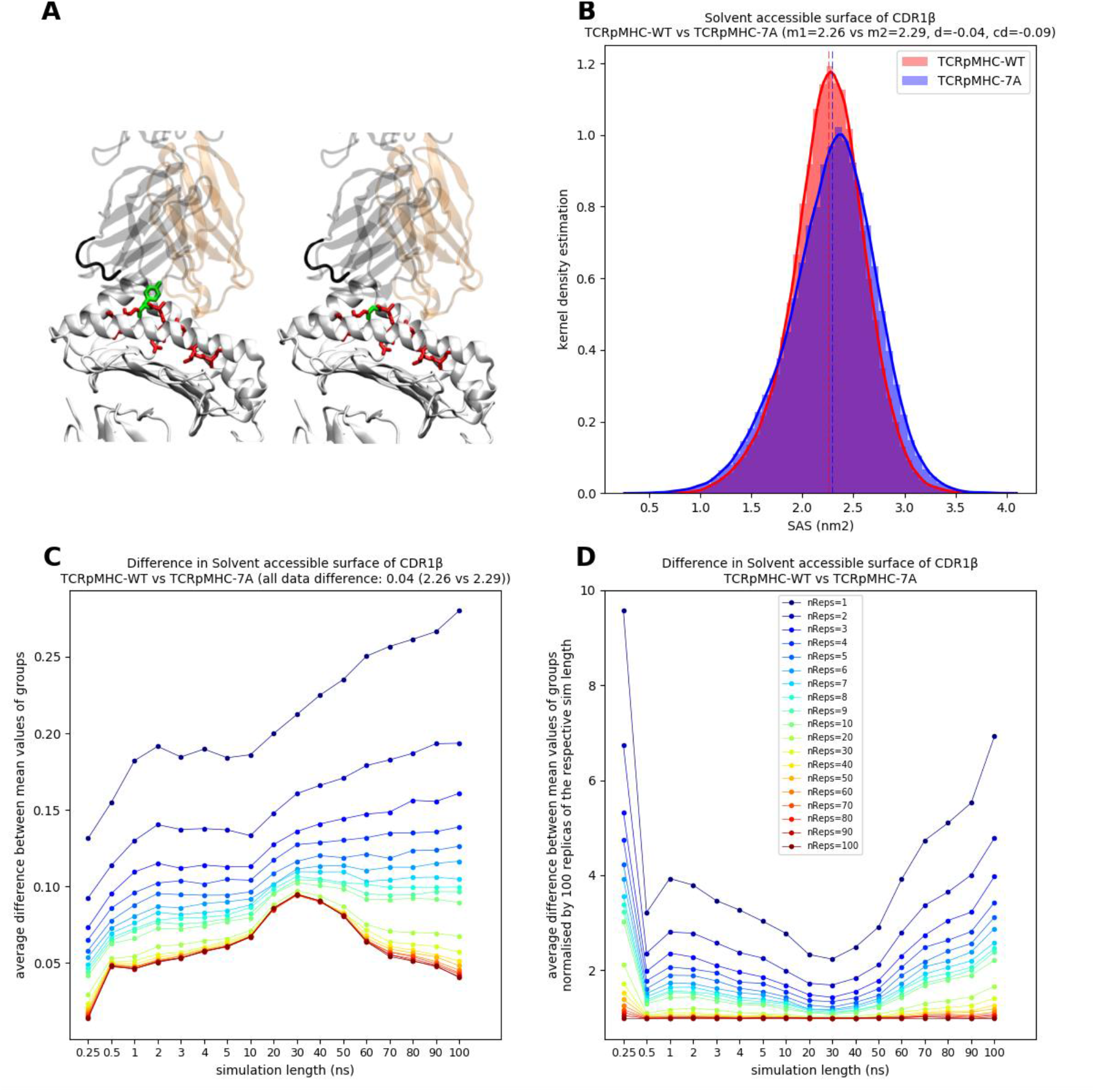
Same as Figure 4 but for the SASA of CDR1β.

**Figure S 18:**
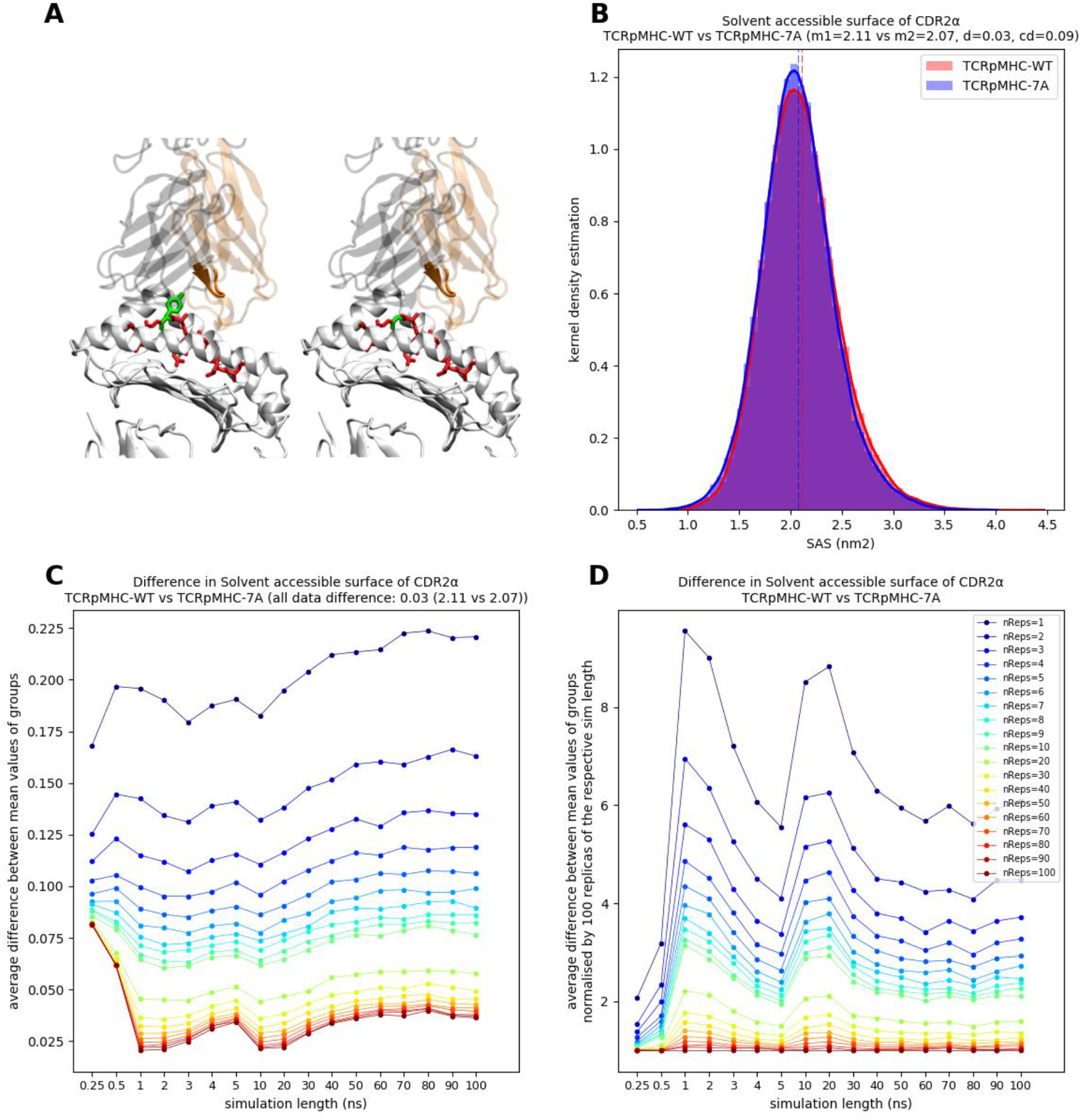
Same as Figure 4 but for the SASA of CDR2α.

**Figure S 19:**
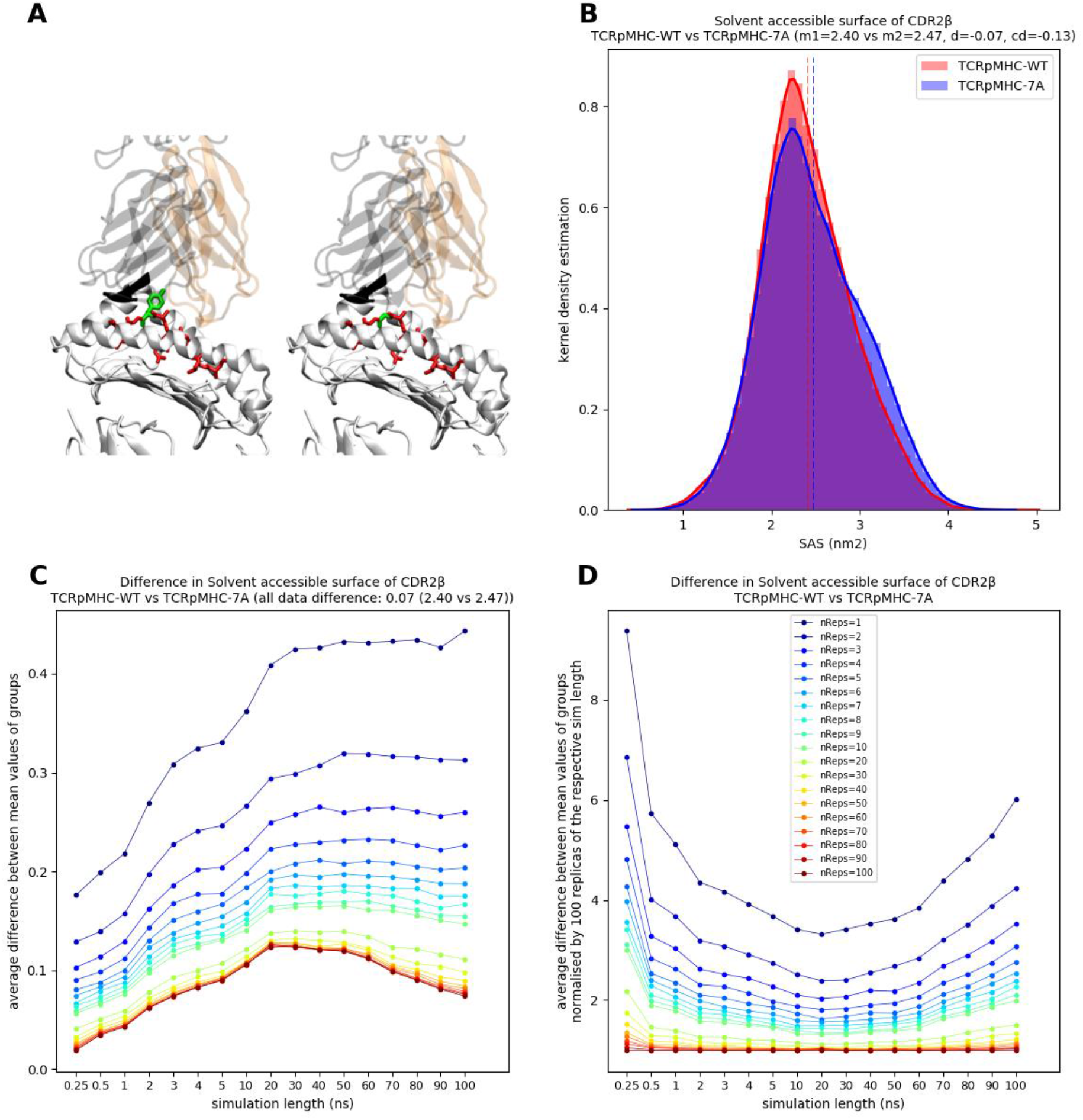
Same as Figure 4 but for the SASA of CDR2β.

**Figure S 20:**
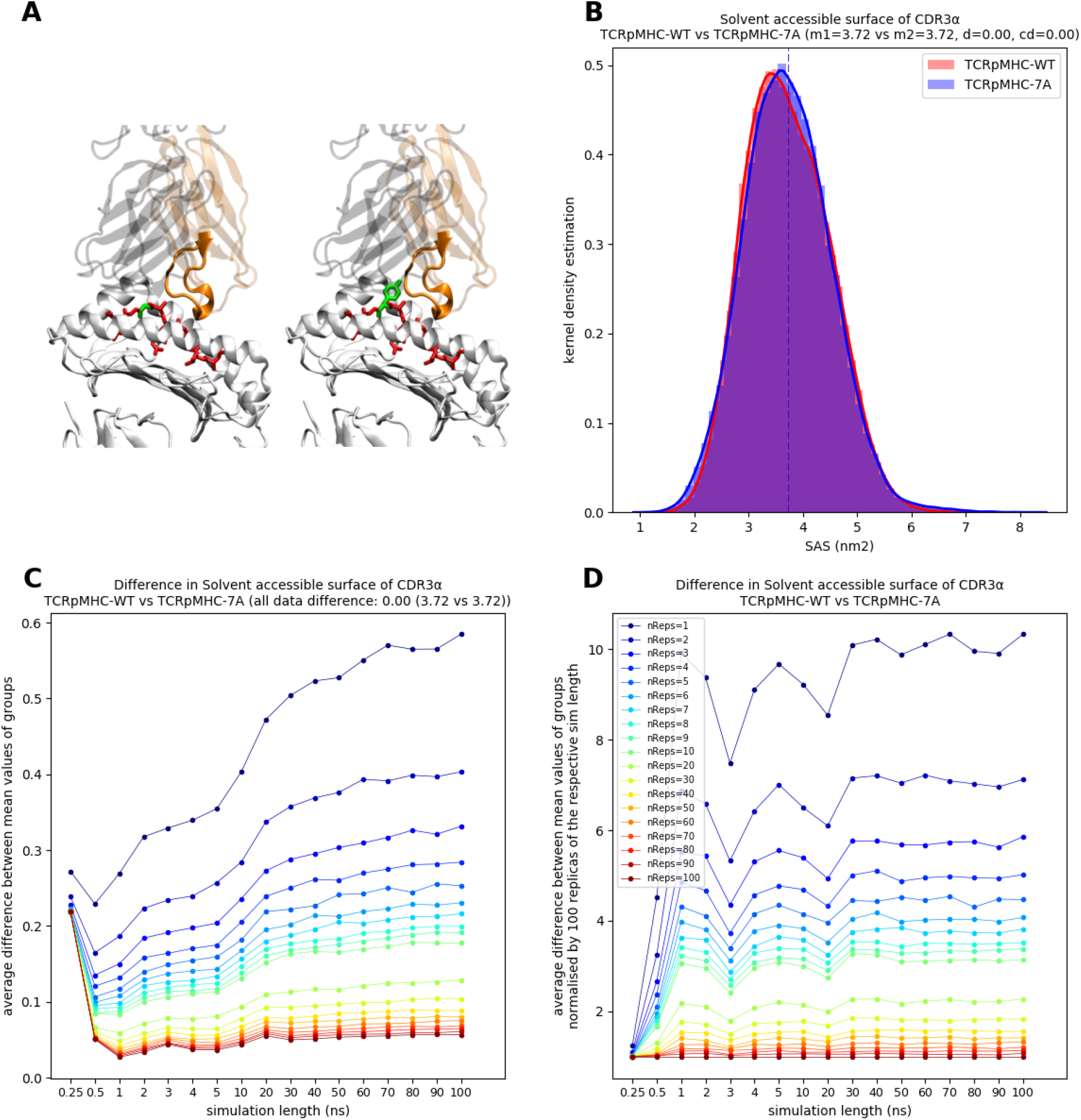
Same as Figure 4 but for the SASA of CDR3α.

**Figure S 21:**
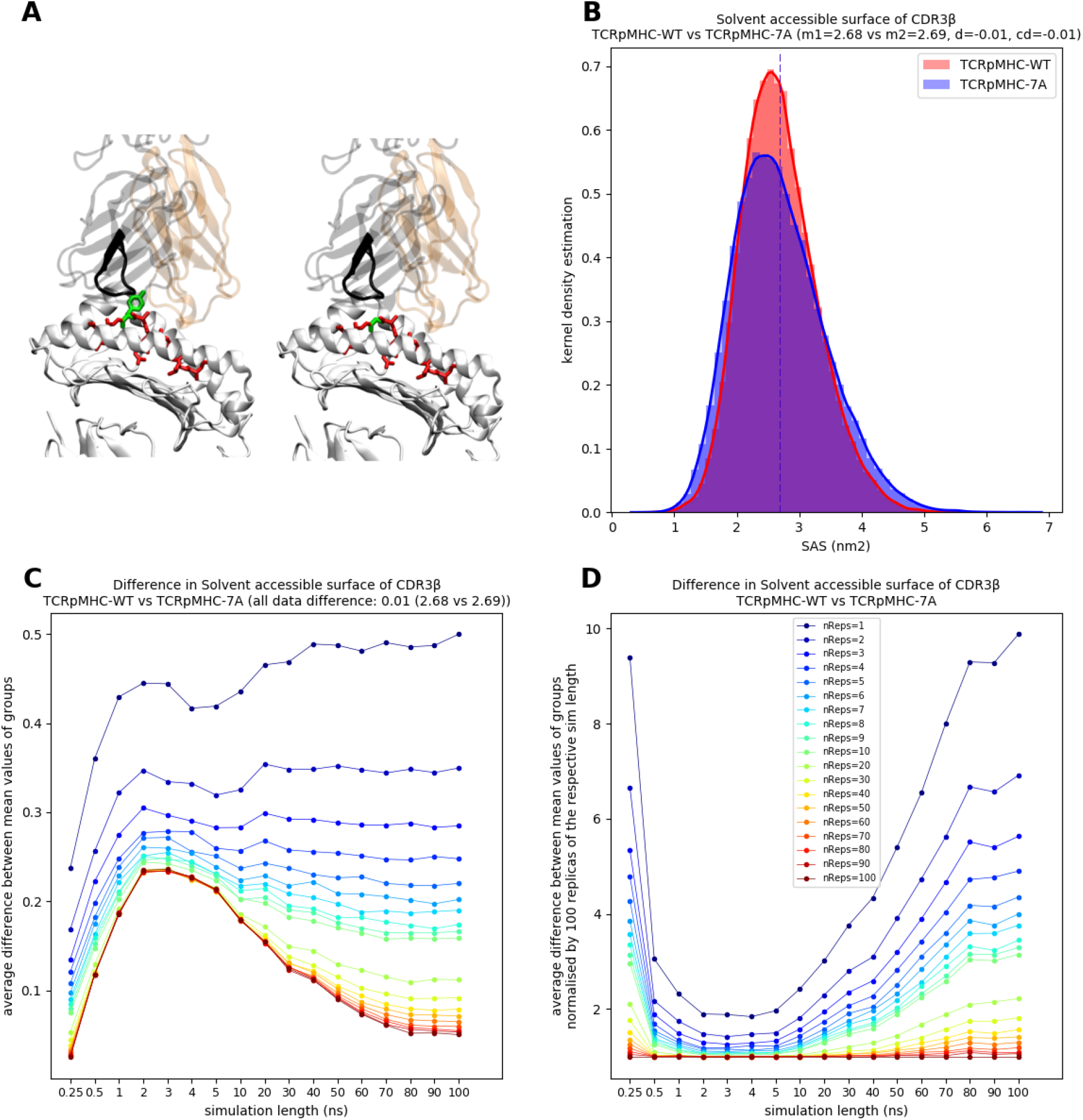
Same as Figure 4 but for the SASA of CDR3β.

